# Structural and mechanistic insights into translation initiation on the enterovirus Type 1 IRES

**DOI:** 10.1101/2025.10.04.680434

**Authors:** Miguel Aracena-Velázquez, Stephen Sukumar Nuthalapati, Jacqueline Hankinson, Ksenia Fominykh, Valeria Lulla, Trevor R. Sweeney, Chris H. Hill

## Abstract

Enteroviruses are a diverse group of pathogens that cause over one billion human infections annually. Upon cell entry, translation of the viral genome is directed by an internal ribosome entry site (IRES) within the 5′ untranslated region. Despite early identification of the Type 1 poliovirus IRES, the structural and mechanistic basis for its activity remains poorly understood due to its size, flexibility and dependence on multiple host cell factors. Here, we reconstitute human translation initiation on a model poliovirus IRES and examine 48S complexes by cryo-electron microscopy. Our structures reveal how IRES domain IVc contacts ribosomal proteins uS19 and uS13, whilst a conserved GNRA tetraloop engages with the initiator tRNA during start-codon recognition. Disruption of these interfaces impairs IRES-dependent translation and viral replication. Together, our results provide new structural and mechanistic insights into initiation on the Type 1 IRES and reveal conserved RNA–RNA interactions critical for enterovirus translation.

## Introduction

The enteroviruses (family *Picornaviridae*) are a group of ubiquitous human and animal pathogens, encompassing poliovirus (PV), enteroviruses, coxsackieviruses, rhinoviruses and echoviruses(Brouwer *et al*, 2021; Simmonds *et al*, 2020). Together, they cause a wide array of diseases, ranging from mild (e.g. respiratory tract infection, self-limiting gastroenteritis) to severe (poliomyelitis, viral meningitis, myocarditis, acute flaccid myelitis) with an estimated global morbidity of more than one billion infections per year (Chang *et al*, 2019; Gonzalez *et al*, 2019; Kohil *et al*, 2021; Li *et al*, 2020; Lulla & Sridhar, 2024; Ndiaye *et al*, 2024; Racaniello, 2006; Suresh *et al*, 2018; Tao *et al*, 2014; Tapparel *et al*, 2013). Upon cell entry, the first molecular event in the infection cycle is translation of the positive-sense, single-stranded RNA genome to produce the viral polyprotein. Gaining control over host ribosomes is therefore an essential event that exemplifies virus-host interaction at a fundamental level – the execution of genetic information.

Most protein-coding messenger RNAs (mRNAs) are translated in a 5′ cap-dependent manner, requiring the coordinated action of multiple eukaryotic initiation factors (eIFs) [reviewed in (Brito Querido *et al*, 2024; Hinnebusch, 2017; Jackson *et al*, 2010; Merrick & Pavitt, 2018)]. The process begins with the assembly of the 43S pre-initiation complex (PIC), comprising the 40S ribosomal subunit bound to eIFs 1, 1A, 3 and the eIF2·GTP·Met-tRNA_i_^Met^ ternary complex. This is recruited to the 5′ cap structure by eIF4F [reviewed in (Gingras *et al*, 1999; Hinnebusch & Lorsch, 2012; Merrick, 2015)], a heterotrimeric complex composed of cap-binding protein eIF4E (Marcotrigiano *et al*, 1997; Sonenberg *et al*, 1978), DEAD-box RNA helicase eIF4A (Caruthers *et al*, 2000; Lawson *et al*, 1989; Rozen *et al*, 1990) and scaffold eIF4G, which also binds to RNA and eIF3 (Kaye *et al*, 2009; Villa *et al*, 2013). The ATP-dependent activity of eIF4A is stimulated by eIF4B and 4G (Andreou & Klostermeier, 2014; Harms *et al*, 2014; Jaramillo *et al*, 1991), facilitating unwinding of secondary structure proximal to the cap, prior to 43S PIC loading. This results in a 48S PIC, which scans along the mRNA in a 5′→3′ direction, searching for a start codon. During scanning, eIF1 and 1A maintain an ‘open’ conformation of the 40S head (Passmore *et al*, 2007) with tRNA in a P^OUT^ position, allowing mRNA to slide past the anticodon until an AUG in good Kozak context (Kozak, 1981) is recognised (Brito Querido *et al*, 2020; Llácer *et al*, 2021; Llácer *et al*, 2015). Upon AUG recognition the tRNA adopts a P_IN_ conformation, the 40S head closes around the mRNA, and eIF1 departs (Hussain *et al*, 2014; Llácer *et al*., 2015; Simonetti *et al*, 2020; Yi *et al*, 2022). Subsequent binding of eIF5 (Llácer *et al*, 2018) activates GTP hydrolysis and P_i_ release by eIF2 (Paulin *et al*, 2001), committing the ribosome to the start codon (Algire *et al*, 2005; Pisarev *et al*, 2006; Unbehaun *et al*, 2004) and triggering further conformational changes (Petrychenko *et al*, 2025). Finally, eIF2·GDP departure and recruitment of eIF5B realign the aminoacyl-acceptor arm of the tRNA for productive 60S joining (Lapointe *et al*, 2022; Pestova *et al*, 2000; Petrychenko *et al*., 2025; Wang *et al*, 2019).

In contrast, viral genome translation is often cap-independent and driven by non-canonical interactions between structured RNA elements and the translation machinery (reviewed in (Lee *et al*, 2017; Mailliot & Martin, 2018; Martínez-Salas *et al*, 2017). Enterovirus translation initiation is facilitated by a ∼450 nt Type 1 internal ribosome entry site (IRES) located within the 5′ untranslated region (UTR). Despite this being the first identified case of internal ribosome entry (Kitamura *et al*, 1981; Pelletier & Sonenberg, 1988), the size and complexity of this element has precluded a detailed structural analysis. Type 1 IRESs share a conserved topology of five structured RNA domains (dII–dVI, **Figure 1A**) and have similar requirements for directing initiation. In addition to the small ribosomal subunit (40S), eIFs 1A, 2, 3, 4A, 4B and the central region of eIF4G [amino acids 736-1115, here denoted as ‘4Gm’, (Pestova *et al*, 1996)] are needed for efficient IRES activity. There was initial controversy regarding which

**Figure 1.**
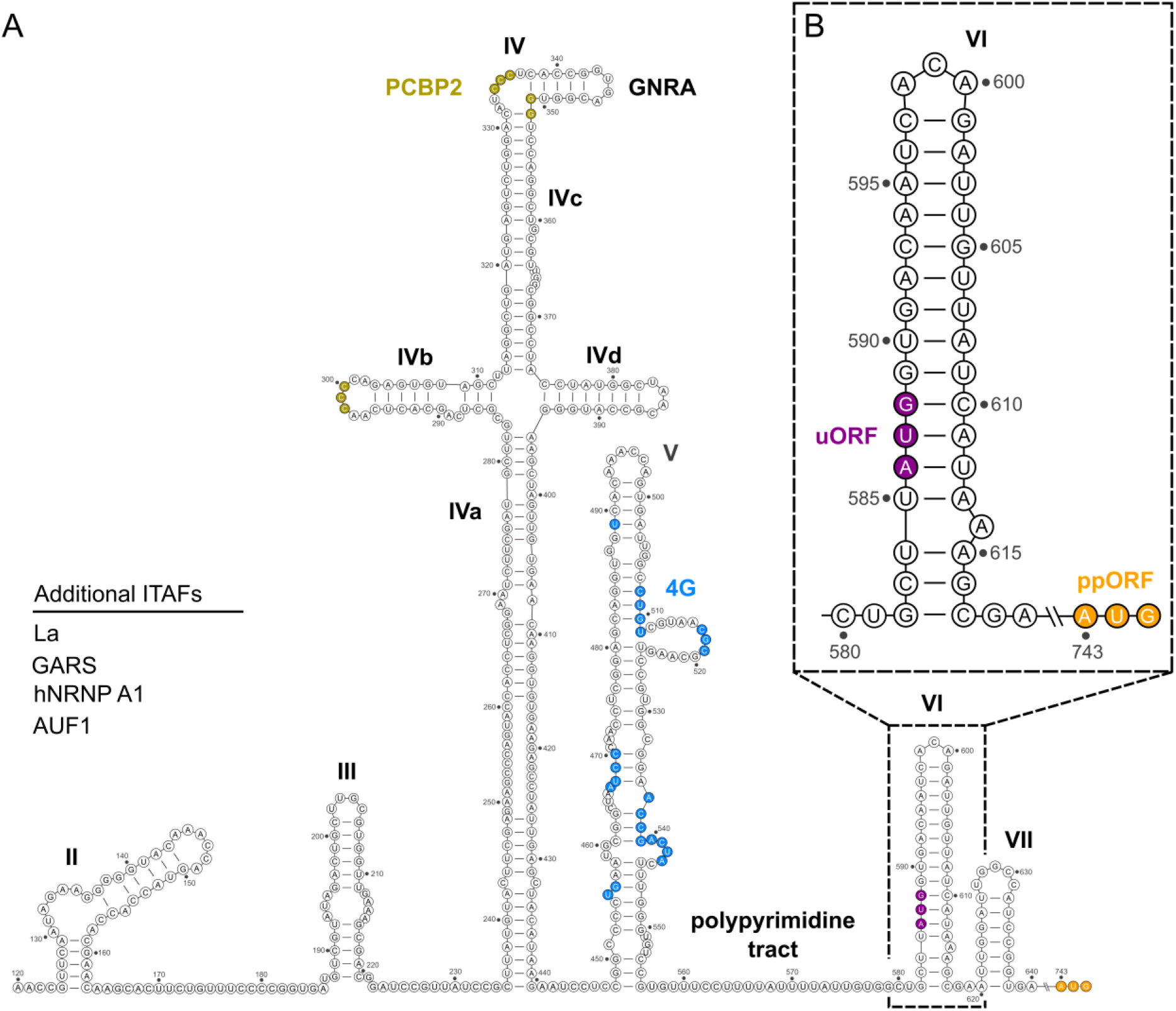
Introducing the enterovirus Type 1 IRES. **A,** Sequence and secondary structure of the poliovirus (PV) IRES. Domains are indicated with bold numerals. Known binding sites for PCBP2 and eIF4G are indicated in beige and blue, respectively. **B,** Close-up view showing details of domain VI. The relative positions of the two AUG codons (uORF AUG586 and ppORF AUG743) are highlighted in purple and orange, respectively.

IRES trans-acting factors (ITAFs) were required – frequently cited ITAFs include glycyl-tRNA synthetase (GARS), La and polypyrimidine tract binding protein (PTB) – although *in vitro* reconstitution experiments have demonstrated that poly-C binding protein (PCBP2) is necessary and sufficient for initiation on several Type 1 IRESs (Andreev *et al*, 2007; Borman *et al*, 1993; Meerovitch *et al*, 1993; Sweeney *et al*, 2014). Notably, eIF4E or an intact eIF4F complex are not required. This allows the virus to selectively inhibit cap-dependent translation of host mRNAs by inducing cleavage of eIF4G through activity of the viral 2A protease (Gradi *et al*, 1998; Lamphear *et al*, 1995).

Several mechanistic features have been identified to date using biochemical and mutagenic approaches, including the binding of eIF4G-4A to dV (de Breyne *et al*, 2009), the recruitment of 43S preinitiation complexes via an eIF4G-eIF3 interaction (Sweeney *et al*., 2014), and the interaction between PCBP2 and dIV (Beckham *et al*, 2020; Sweeney *et al*., 2014). Furthermore, enteroviruses contain an alternate upstream open reading frame (uORF) encoding a protein necessary for viral infection in gut epithelial cells (Lulla *et al*, 2019). The AUG codon directing translation of this product is located within dVI, and this upstream AUG is also essential for initiation at the main polyprotein open reading frame (ppORF) (Iizuka *et al*, 1991; Kaminski *et al*, 2010; Meerovitch *et al*, 1991; Nicholson *et al*, 1991; Pestova *et al*, 1994). However, mechanisms of uORF versus ppORF ribosome commitment are not understood, and the occlusion of uORF AUG within dVI implies that unwinding of dVI must occur during productive initiation.

Although evolutionarily distinct, Type 1 IRESs share some common structural motifs with the similarly large Type 2 IRESs found, for example, in encephalomyocarditis virus (EMCV) and foot-and-mouth disease virus (FMDV). These include a conserved polypyrimidine tract immediately upstream of the AUG start codon nearest the 3′ border of the IRES, and a GNRA-loop structure at the apex of the largest IRES domain (equivalent to Type 1 dIV, **Figure 1A**), both of which are necessary for IRES activity (Fernández-Miragall & Martínez-Salas, 2003; Fernández *et al*, 2013; Fernández *et al*, 2011; López de Quinto & Martínez-Salas, 1997; Nateri *et al*, 2000; Robertson *et al*, 1999). Nevertheless, we lack an understanding of how this complicated molecular machinery is organised in three dimensions, and the network of IRES-ribosome interactions remains largely uncharacterised. In addition, the mechanism for activation by any ITAF, including PCBP2, has not been determined.

To address this, here we combine cryogenic electron microscopy (cryo-EM), translation assays, virological assays and RNA structure probing to elucidate the mechanistic basis of translation initiation on the Type 1 IRES from poliovirus (PV) and related enteroviruses. Our cryo-EM structures of human 48S initiation complexes reveal a network of interactions between the IRES and the translation machinery. The conserved GNRA loop at the apex of dIV binds to the initiator tRNA in the ribosomal P-site, and the dIV RNA is further stabilised by direct contacts with uS13 and uS19 on the 40S ribosomal subunit. We next demonstrate that disruption of these interactions has a severe impact on protein synthesis and replication in closely related coxsackievirus CVA13. Together, our work provides new insights into the structure and function of a critical viral RNA element that has remained enigmatic since its discovery nearly 40 years ago.

## Results

### Visualising initiation on the poliovirus Type 1 IRES

While progress has been made in visualising multiple stages of initiation on smaller CrPV-like IRESs (Abaeva *et al*, 2020; Acosta-Reyes *et al*, 2019; Fernández *et al*, 2014; Muhs *et al*, 2015; Murray *et al*, 2016; Pisareva *et al*, 2018; Spahn *et al*, 2004) and HCV-like IRESs (Brown *et al*, 2022; Hashem *et al*, 2013; Quade *et al*, 2015; Yamamoto *et al*, 2015; Yamamoto *et al*, 2014; Yokoyama *et al*, 2019), structural information on the organisation of larger, more complicated IRESs is lacking, and the network of IRES-ribosome interactions that promote translation remains largely uncharacterised. We therefore decided to investigate translation initiation on the PV Type 1 IRES using cryogenic electron microscopy (cryo-EM). Initiation occurs at two alternate AUG codons on the PV IRES: AUG_586_ at the 3′ border of the minimal IRES region and AUG_743_ at the start of the long polyprotein open reading frame (**Figure 1B**). AUG_586_ dependent translation was recently confirmed to enhance viral replication in intestinal organoid cultures (Lulla *et al*., 2019). To reduce heterogeneity for structural work, we used a PV IRES mutant, “PV-AUG_586_-good”, in which AUG_586_ has been placed in an optimal Kozak sequence to promote efficient 48S complex formation (Pestova *et al*., 1994) and AUG_743_ is absent (**Figure 2A**).

**Figure 2.**
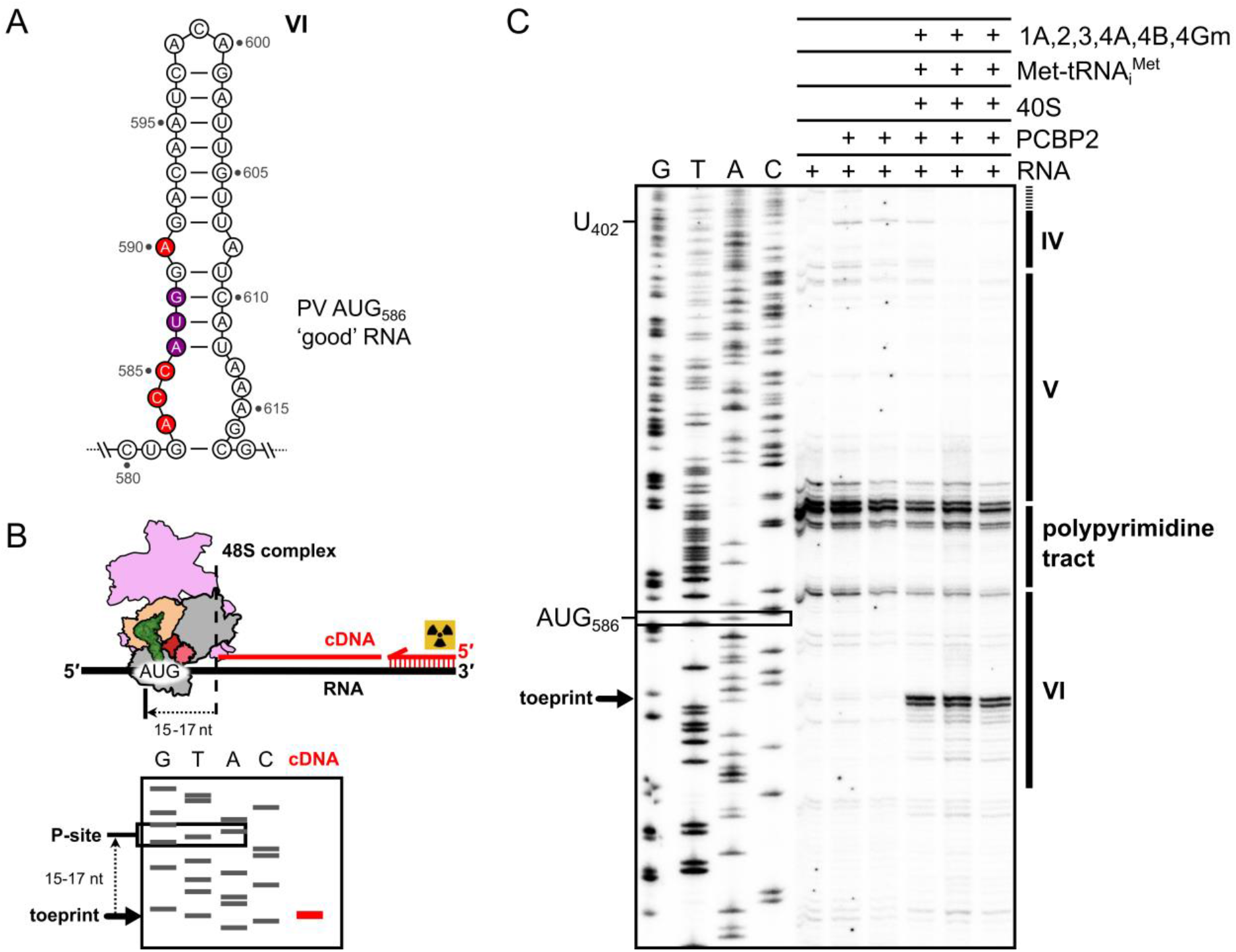
Reconstitution of human 48S initiation complexes on the poliovirus Type 1 IRES. **A,** Details of genetic changes (red) to the model PV-AUG586-good RNA. The uORF AUG present in domain VI is placed in an ideal Kozak sequence context (Sweeney et al., 2014). **B,** Initiation is monitored by a termination of primer extension assay (toeprinting). If 48S complexes form, the 3′ edge of the ribosome arrests the reverse transcriptase 15-17 nt downstream of the +1 position in the P-site. The exact location of the P-site is inferred from the adjacent Sanger sequencing reaction, run on the same gel using the same primer. **C,** Analysis of primer extension assay by Urea-PAGE and autoradiography, demonstrating human 48S complex formation at AUG586 on the PV-AUG586-good IRES. The expected position of toeprints linked to initiation at AUG586 is indicated (black arrow). IRES domains corresponding to the displayed sequencing ladder are indicated on the right. Three independent reactions are shown alongside negative controls omitting ribosomes and initiation factors.

First, we reconstituted translation initiation *in vitro* using purified human ribosomes and translation initiation factors (**Figure EV1**). <u>T</u>ermination <u>o</u>f primer <u>e</u>xtension (<u>toe</u>printing) assays confirmed efficient 48S complex assembly at AUG_586_ in the presence of 40S, Met-tRNA_i_^Met^, eIFs 1A, 2, 3, 4A, 4B, 4Gm and ITAF PCBP2 (**Figure 2B,C**). To verify that our 48S complexes were forming via an authentic pathway, we performed factor drop-out experiments (**Figure EV2, Appendix Figure S1A**). Omission of 40S, Met-tRNA_i_^Met^, eIFs 2, 3 or 4A disabled 48S complex formation at AUG_586_ (<10% control, background levels). Omission of 4Gm had severe effects (∼30% control) whereas omission of eIF4B, eIF1A or PCBP2 resulted in reduced efficiency of 48S assembly (∼50-70% control, variable between experiments). This is consistent with previous observations *in vitro* (Sweeney *et al*., 2014) suggesting that our 48S complexes were assembling on the PV-AUG_586_-good IRES via a mechanistically relevant pathway.

Having confirmed 48S assembly at AUG_586_, we next reconstituted complexes for structural analysis by mixing 40S subunits, Met-tRNA_i_^Met^, eIFs 1A, 2, 3, 4A, 4B, 4Gm, PCBP2, PV-AUG_586_-good RNA, ATP and GTP. To capture early stages of initiation (prior to GTP hydrolysis by eIF2, or eIF2·GDP departure) we omitted eIF1, eIF5 and eIF5B from the reconstitution. After mixing the components, we incubated complexes at 37°C for 15 min, prepared grids without crosslinking and collected cryo-EM data (**Figure 3A**). Following acquisition of 26,797 micrographs, extensive particle classification yielded several distinct structures including 40S, 40S-eIF3-eIF1A, an ‘open’ P_OUT_ 48S and ‘closed’ P_IN_ 48S complexes at AUG_586_, with and without eIF3 (**Figure 3B, Figure EV3, Figure EV4, Appendix Figure S2-S5, Appendix Table S1**).

**Figure 3.**
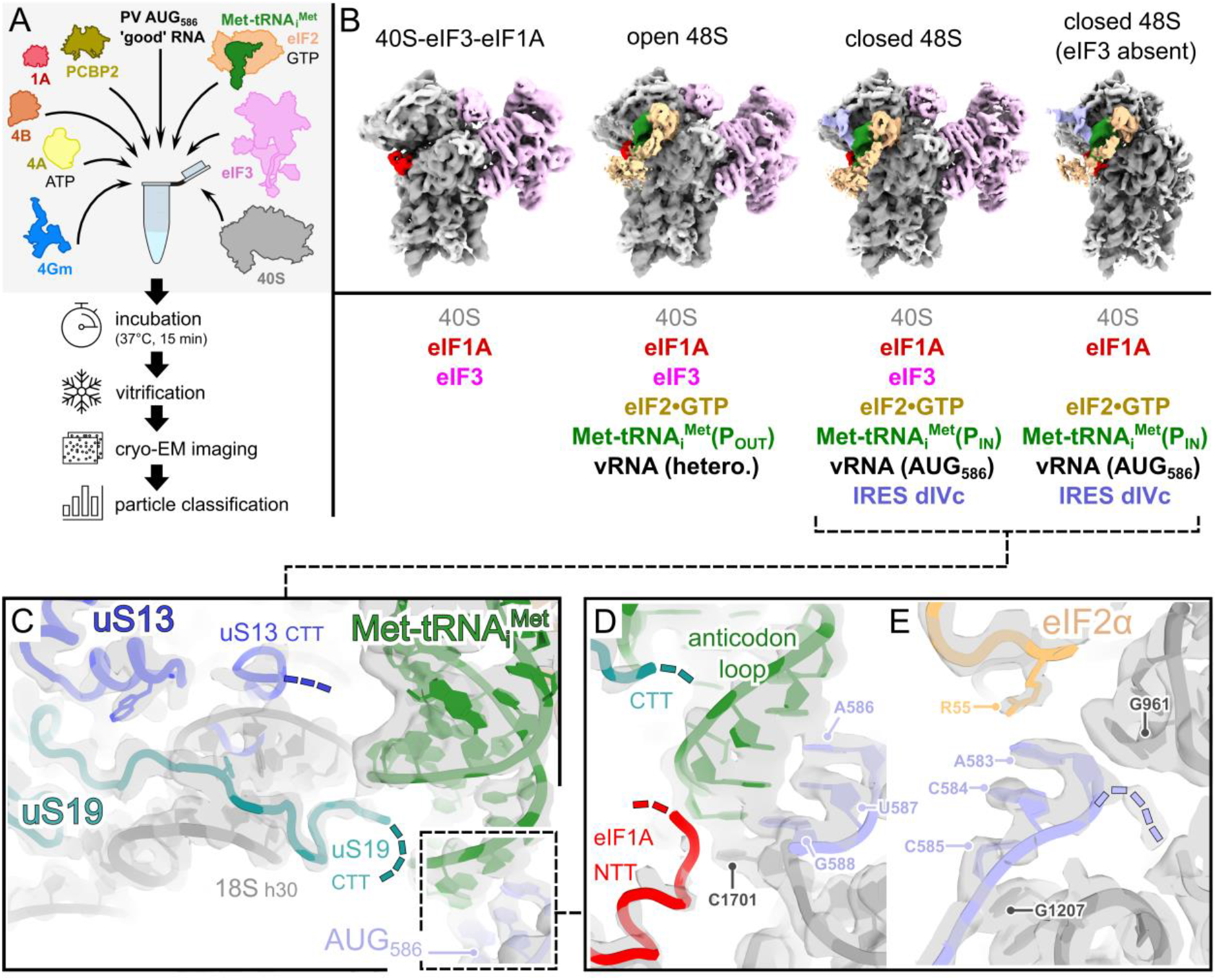
cryo-EM structures of human 48S complexes during PV initiation. **A,** Schematic diagram illustrating our workflow for structural investigation of 48S initiation complexes. **B,** Particle classification (see Figures EV3 and EV4) allowed for separation of four major states. Cryo-EM maps from 3D refinement of key classification intermediates are shown (downsampled pixel size of 3.36 Å/px, contoured at 3.0–5.0 σ) and colour-coded based on the identity of components, as listed below. **C,** Details of the mRNA channel in ‘closed’ 48S complexes. Local resolution weighted cryo-EM maps from structure 2 are displayed (grey), contoured at 5.0 σ (estimated local resolution 3.0 - 3.5 Å). C-terminal tails (CTT) of uS19 (turquoise) and uS13 (blue) extend towards the initiator tRNA (green) but are not fully ordered as observed in post-GTP hydrolysis structures (Petrychenko et al., 2025). **D,** Details of codon-anticodon interaction in the P-site. Cryo-EM maps (grey) are displayed as in C. Density for the N-terminal tail of eIF1A (red) is observed, consistent with recognition of AUG586 in dVI of the PV IRES. **E.** Details of Kozak residues in the E-site. Cryo-EM maps (grey) are displayed as in C, contoured at 4.0 σ. Contacts with 18S G1207 and G961 are observed at -1 and -3 positions, respectively. R55 from eIF2α also contacts A583 at -3.

In our ‘open’ P_OUT_ 48S map (**structure 1**) the relative positioning of the 40S head and body is consistent with an open conformation (Llácer *et al*., 2015) as determined by the distance between h34 and h18 of the 18S rRNA at the mRNA latch region (**Appendix Figure S6A**). This conformation is nearly identical to another human scanning 48S complex (Brito Querido *et al*., 2020) (Data ref: PDB 6ZMW, 2020; **Figure 3B**) but differs from other ‘open’ 48S structures (Petrychenko *et al*., 2025; Yi *et al*., 2022) in the amount of head rotation (Data ref: PDB 8PJ1, 2024; Data ref: PDB 7QP6, 2022; **Appendix Figure S6A**). We also observe differences in the positioning of eIF2 and tRNA. In the absence of eIF1 in our reconstitutions, eIF2β moves ∼16 Å towards 18S A1058 at the eIF1 binding site (**Appendix Figure S6B**). As a result, the initiator tRNA and eIF2 are tilted towards the E-site, leading to shifts of up to ∼14 Å in the position of the tRNA aminoacyl-acceptor arm, and ∼5 Å in the position of the anticodon (**Appendix Figure S6C**). The 40S mRNA channel contains weak heterogeneous density, but no codon-anticodon interactions are visible. It is unclear whether this structure represents a vRNA loading intermediate, or whether the observed differences in tRNA and eIF2 position are caused by omission of eIF1.

In contrast, our ‘closed’ P_IN_ 48S structures exhibit unambiguous codon-anticodon base pairing between initiator tRNA and AUG_586_ mRNA in the P-site, a closed 40S head position (Llácer *et al*., 2015), and ordered density for eIF1A N-terminal tails (NTT), and uS13 and uS19 C-terminal tails (CTT), respectively (**Figure 3C-E**). To enable inspection of AUG_586_, dVI is completely unwound, with no structured RNA elements occupying either A- or E-sites. The conformation of eIF2 in our closed 48S complexes most closely resembles that prior to GTP hydrolysis (Petrychenko *et al*., 2025), and we observe weak density for the eIF2β subunit. eIF3 is present in ∼40% of the P_IN_ closed particles at AUG_586_, but the flexible peripheral subunits eIF3b, 3g and 3i are not well-resolved. eIF3j is also absent, consistent with its dissociation from the 43S PIC upon mRNA binding (Fraser *et al*, 2007; Sokabe & Fraser, 2017). The N-terminal domain (NTD) of eIF3c (1-160) is not bound to the 40S surface in our maps, possibly due to the lack of stabilising interactions with eIF1 (Yi *et al*., 2022) which was omitted from our reconstitutions. This is consistent with previous observations of late-stage mammalian 48S complexes after eIF1 dissociation (Simonetti *et al*., 2020). We did not observe ordered density for eIF4Gm or 4A in any of our maps.

### IRES domain IVc forms a network of contacts with the translation machinery

In our ‘closed’ 48S complexes, we observed additional density bridging the 40S subunit head with the initiator tRNA (**Figure 3B** and **Figure 4A,B; structure 2**). Whilst the decoding centre of the ribosome is well-resolved, flexibility at the periphery restricted the local resolution of the map in this region (∼3.5-8.0 Å). Nevertheless, it was sufficient to model the RNA backbone, and to determine the lengths and relative orientations of helical components. Comparison with potential IRES elements present in our construct allowed us to identify this density as domain IVc (dIVc) comprising nucleotides 313-374 of the PV IRES (**Figure 4B**). Based on the principal modes of flexibility revealed by focused classification, we propose the further subdivision of dIVc into three helical segments oriented at ∼90° to each other (**Figure 5A**). dIVc_1_ (313-319; 368-374) is the most flexible, and we were able to separate two conformations for this segment (**structures 3 and 4**). It extends from the ribosome-distal dIVc-IVb-IVd junction towards the ribosome surface, before a sharp turn at the uS19 contact site. Beyond this, dIVc_2_ (323-331; 352-360) extends laterally across the surface of uS13, leading to dIVc_3_ (332-351): a C-rich loop connected to a short helix (5 bp), capped by an apical GNRA tetraloop. No density for any other IRES domain was observed, suggesting that dII, dIII and dV are flexible with respect to dIV, and in the state observed here, these domains lack equivalent stabilising interactions with the ribosome surface.

**Figure 4.**
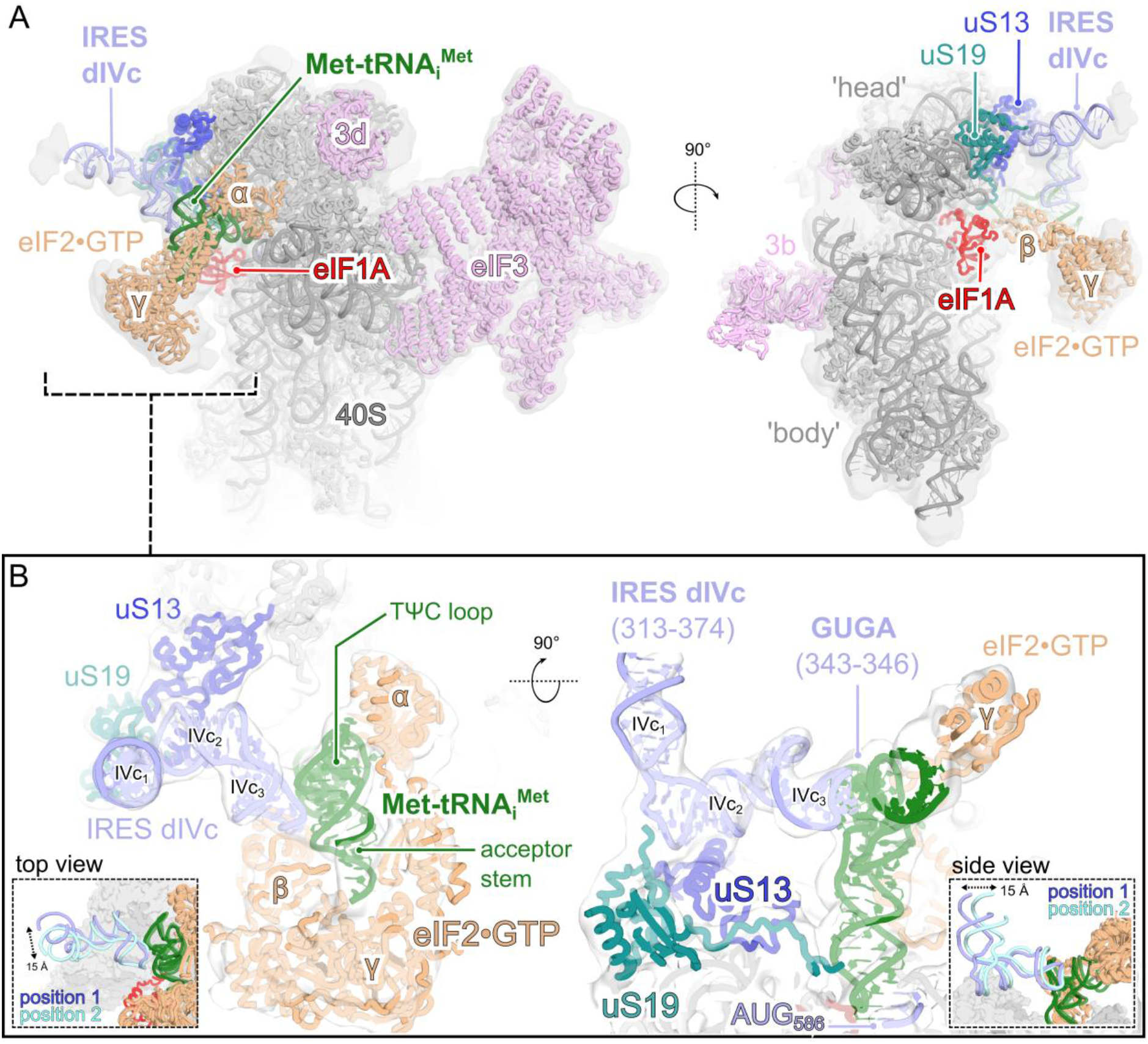
IRES domain IVc bridges the 40S head and the initiator tRNA in ‘closed’ 48S complexes. **A,** Overall architecture of the ‘closed’ 48S complex at PV AUG586. Subunits and components are colour-coded as in Figure 3. Cryo-EM maps from 3D refinement of structure 2 (grey) are displayed (lowpass filtered to 10 Å and contoured at 1.4 σ) to show occupancy of flexible components. Observation of eIF2β suggests a state pre-GTP hydrolysis, consistent with the omission of eIF5. **B,** Close-up view of contacts between IRES domain IVc and the translation machinery. Domain IVc bridges the 40S head with the initiator tRNA through contacts with uS19, uS13 and initiator tRNA. (Inset) The stem region of dIVc beyond the contacts with the ribosome surface (313-322; 361-374) exhibits flexibility, limiting local resolution. Focused classification was able to resolve two conformations (structures 3 and 4), related by a helical displacement of ∼15 Å.

**Figure 5.**
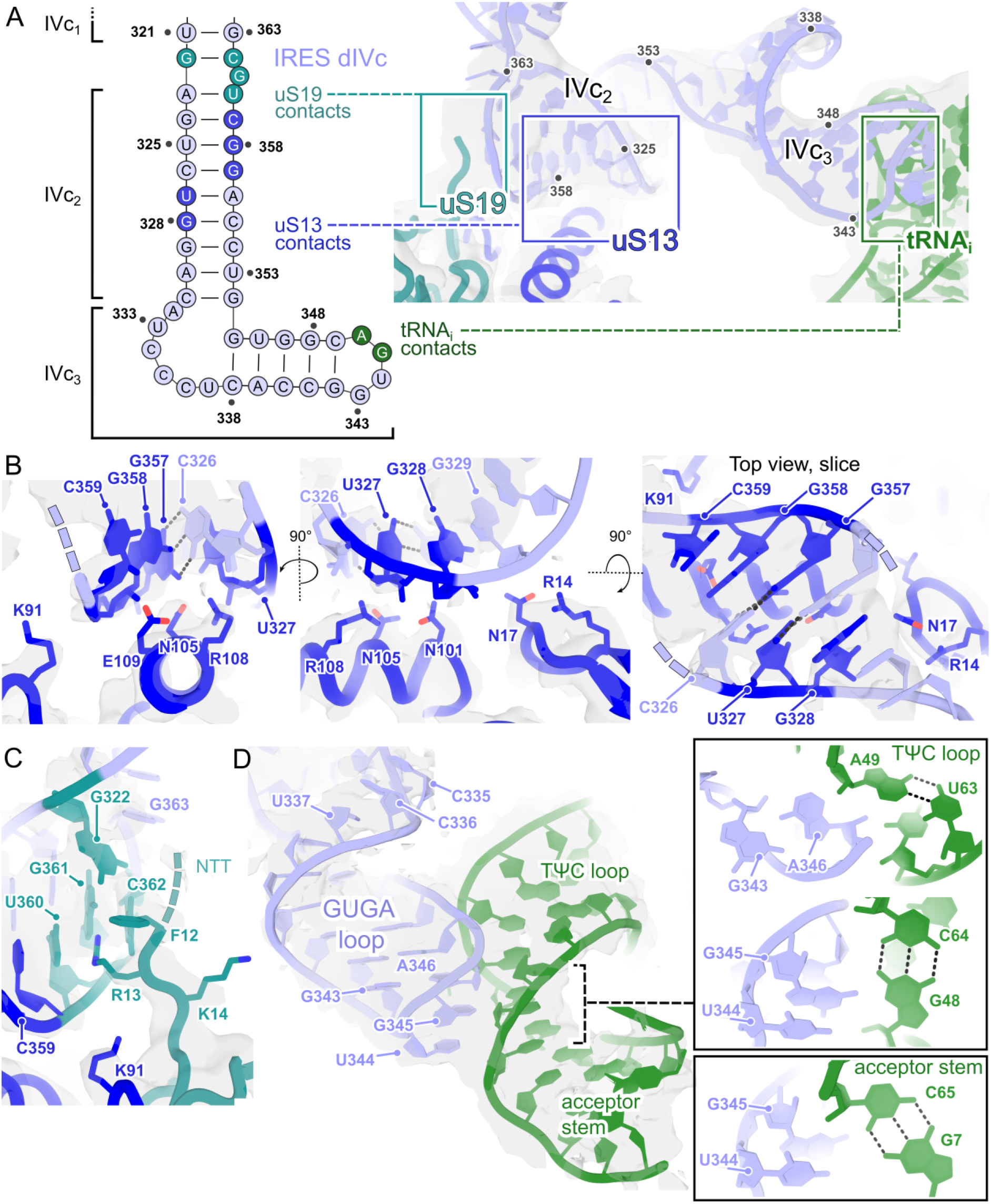
Details of contacts between IRES domain IVc and the translation machinery. **A,** (Left) Domain IVc sequence diagram, summarising key contacts. Nucleotides are colour-coded to indicate interaction with uS13 (blue), uS19 (turquoise) or initiator tRNA (green). (Right) Areas of interest are indicated on the model, colour coded as above. Local resolution weighted cryo-EM maps from structure 2 are displayed (grey), contoured at 2.8 σ. The resolution of the displayed region is estimated to be ∼3.5-8.0 Å. **B,** Close-up view of interaction with uS13 (blue), shown in three orthogonal views. Amino acid side-chains at the interface are shown as sticks. Local resolution weighted cryo-EM maps from structure 2 are displayed (grey), contoured at 4.5 σ. Nucleotides within 4.0 Å of labelled uS13 residues are highlighted (blue). The interaction surface is centred around the G357-C326 base pair. **C,** Close-up view of interaction with the N-terminal tail (NTT) of uS19 (turquoise). Amino acid side-chains at the interface are shown as sticks. Local resolution weighted cryo-EM maps from structure 2 are displayed (grey), contoured at 4.5 σ. Nucleotides within 4.0 Å of labelled uS19 residues are highlighted (turquoise). **D,** Close-up view of interaction with initiator tRNA (green). Cryo-EM maps from 3D refinement of structure 2 are displayed (grey), contoured at 3.5 σ. The apical GUGA tetraloop docks in the minor groove of the helical junction between the TѰC loop and the acceptor stem during AUG recognition. (Inset) G345 and A346 are within 4.0 Å of tRNAi base pairs A49-U63, G48-C64 (TѰC loop) and G7-C65 (acceptor stem).

The dIVc interface with the 40S subunit occurs at a contiguous surface formed by proteins uS13 and uS19. Helix α5 of uS13 acts as a docking platform for the minor groove of the dIVc_2_ helix, with N101, R108 and E109 forming polar contacts with the ribose phosphate backbone (**Figure 5B**). N105 is positioned adjacent to the G357-C326 base pair and may directly interact with the guanine base. In contrast, the flexible N-terminus of uS19 packs against the junction between dIVc_1_ and dIVc_2_ segments, comprising U360, G361, C362 and G322. uS19 residues F12, R13 and K14 are ordered (**Figure 5C**), however the disordered basic amino acid sequence preceding this (MAEVEQKKKRT) may also contribute to dIVc_1_ RNA backbone interactions distal to the ribosome.

The GNRA tetraloop in dIVc_3_ (GUGA in PV; 343-346) is highly conserved and essential for initiation on both Type 1 and Type 2 IRESs (Fernández-Miragall & Martínez-Salas, 2003; Fernández *et al*., 2013; Fernández *et al*., 2011; López de Quinto & Martínez-Salas, 1997; Nateri *et al*., 2000; Robertson *et al*., 1999). In our structures, this tetraloop directly contacts the initiator tRNA via insertion of G345 and A346 into the minor groove at the junction between the tRNA acceptor stem and the TѰC loop (**Figure 5D**). G345 and A346 are located <4.0 Å from tRNA_i_ base pairs A49-U63, G48-C64 (TѰC loop) and G7-C65 (acceptor stem). Whilst G343 does not directly interact with tRNA_i_, it may form a non-canonical base pair with A346, stabilising its orientation for productive interaction with the tRNA_i_ (**Figure 5D**). Interestingly, the GNRA-tRNA_i_ interaction occurs exclusively with P_IN_ tRNA_i_ during AUG recognition: this contact is not visible in our ‘open’ 48S complexes, suggesting an incompatibility with the P_OUT_ tRNA_i_ conformation in which the acceptor arm of the tRNA_i_ is ∼4 Å further away from N105 on uS13 (**Appendix Figure S7**). However, very weak density can be observed adjacent to uS19 and uS13 in our ‘open’ 48S complex (**Appendix Figure S7A**). Whilst focused classification was unable to resolve clear density for dIVc_1_ and dIVc_2_ segments in ‘open’ 48S complexes, dIVc may nevertheless contact this site prior to start codon recognition.

Following this logic, we next asked whether the dIVc-uS19-uS13 interactions might independently occur at earlier stages of initiation. Despite exhaustive focused classification, we were unable to detect these interactions in subsets of particles corresponding to 40S or 40S-eIF3-eIF1A. This is consistent with the small buried surface area of the interaction (∼587 Å^2^, **Appendix Figure S8A,B**) and suggests that IRES-mediated initiation proceeds via a multivalent, avidity-driven mechanism. Previously-reported interactions such as dV-eIF4G (de Breyne *et al*., 2009) and eIF4G-eIF3 (Sweeney *et al*., 2014), while not observed here, may nevertheless contribute to overall 48S complex stability.

### The domain IVc contacts are incompatible with later stages of initiation

In our experiments, we artificially trapped early initiation events by omitting eIF5 to delay GTP hydrolysis by eIF2. Later stages of initiation, post eIF2 GTP-hydrolysis, are accompanied by coordinated movements of the tRNA_i_ prior to large subunit joining, associated with the departure of eIF2ꞵ from the decoding centre, the arrival of eIF5B and the dissociation of eIF2 (Petrychenko *et al*., 2025). These movements result in a ∼12 Å net migration of the tRNA_i_ acceptor stem from its initial position in **structure 2** and would break the contact with the GNRA tetraloop in dIVc_3_ (Data ref: PDB 8PJ2, 2024; Data ref: PDB 8PJ3, 2024; Data ref: PDB 8PJ4, 2024; Data ref: PDB 8PJ5, 2024; **Figure 6A**). Moreover, comparison of our structure with human 80S initiation complexes (Holm *et al*, 2023) reveals that the position of dIVc observed here is incompatible with 60S joining (Data ref: PDB 8G5Y, 2023; **Figure 6B**). Whilst intersubunit bridge B1a (involving 60S H38) could form adjacent to dIVc_1_, the B1b/c bridge (60S H84, uL5) would clash with dIVc_3_. Therefore, without large conformational changes in dIVc, the contacts observed here are likely unique to early initiation complexes.

**Figure 6.**
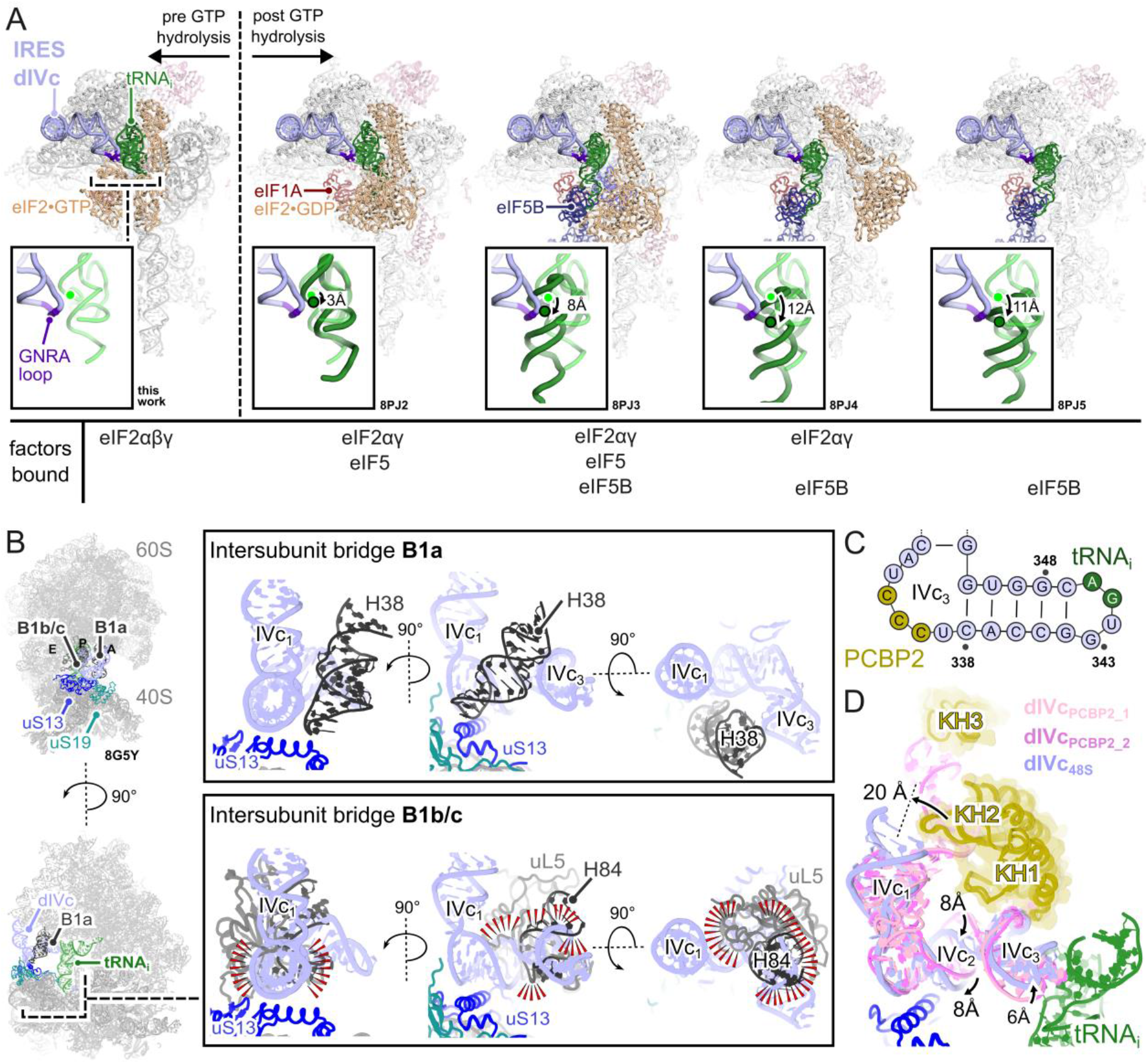
The position of dIVc is incompatible with later stages of initiation and large subunit joining. **A,** Structural superposition of ‘closed’ 48S complexes observed here (structure 2, left) including the PV IRES dIVc, with later human 48S intermediates post GTP hydrolysis Data ref: PDB 8PJ2, 2024, Data ref: PDB 8PJ3, 2024, Data ref: PDB 8PJ4, 2024, Data ref: PDB 8PJ5, 2024 (Petrychenko et al., 2025). Close-up views indicate changes in tRNA position in each state (dark green) compared to the position observed here (light green). Measurements indicate movement relative to A49 in the TѰC loop. The dIVc-tRNA contacts would not persist upon eIF5B binding due to movements of tRNA prior to 60S joining. **B,** Structural superposition of ‘closed’ 48S complex (this work, structure 2) with human 80S initiation complexes Data ref: PDB 8G5Y, 2023 (Holm et al., 2023). The dIVc3 (lilac) would clash (red wedges) with the B1b/c intersubunit bridge comprising uL5 and H84 from the 60S subunit. Formation of the B1a bridge (H38-uS13) may still be possible next to the dIVc-uS19 interface. **C,** Diagram indicating the known PCBP2 binding (brown) site on the PV sequence in the tip of dIVc. **D,** Structural superposition of dIVc observed in 48S complexes (this work, structure 2, lilac) with two models of dIV (pink) bound to PCBP2 (brown) (Beckham et al., 2020). PCBP2 binding would not clash with eIF2, tRNA or the ribosome. Significant displacements between equivalent backbone atoms upon ribosome binding are indicated by black arrows.

The C-rich loop in dIVc_3_ has previously been identified as a binding site for the KH1 domain of PCBP2 (Asnani *et al*, 2016; Beckham *et al*., 2020; Sweeney *et al*., 2014) (**Figure 6C**) but we did not observe density for PCBP2 in any of our structures. We therefore asked whether the conformation of dIVc observed here was compatible with previous cryo-EM and SAXS reconstructions of the PV dIV-PCBP2 complex (Jaqueline Wilce, personal communication). Whilst structural superposition revealed high local similarity between the dIVc_2_ (0.85 Å RMSD) and dIVc_3_ (1.7 Å RMSD) segments, global movements of these segments are necessary to make contacts with uS13 and tRNA_i_, respectively (**Figure 6D**). Whilst PCBP2 would not clash with the tRNA or eIF2, it is unclear to what extent it would be able to efficiently interact with the C-rich loop in dIVc_3_ given the altered dIVc conformation observed here. KH1 and KH2 domains in PCBP2 form a back-to-back dimer (Du *et al*, 2008) and, in a previous model of the dIV-PCBP2 complex, KH2 was proposed to interact with nucleotides 286-288 in the dIVc-IVb-IVd junction, thereby pulling dIVc_1_ towards the GNRA tetraloop at the tip of dIVc_3_ (Beckham *et al*., 2020). In our structures, this restraint does not exist, and we observe a ∼20 Å shift of the dIVc_1_ helical axis away from this position (**Figure 6D**). Whilst PCBP2 binding may be required to initially ‘prime’ dIVc into a structure permissive for ribosome encounter, it is clear from our structures that the contacts with uS13, uS19 and tRNA_i_ can persist in the absence of PCBP2.

To explore this further, we performed RNA structure probing experiments of the PV IRES using the SHAPE (Selective 2′-Hydroxyl Acylation analysed by Primer Extension) technique. In this method, *in vitro* transcribed RNA is modified by incubation with N-methyl isatoic anhydride (NMIA) which reacts with the 2′-hydroxyl group of RNA. This modification is detected as a premature stop in a reverse transcription reaction, with unpaired bases characteristic of unstructured RNA being more reactive than those trapped in secondary structure elements. **Figure 7** shows that the SHAPE reactivity profile generally agrees with the predicted secondary structure depicted in **Figure 1A** based on base-pairing potential. An interesting deviation occurs between dIVc_1_ and dIVc_2_ where high SHAPE reactivity is detected for predicted base-paired nucleotides (**Figure 7A**, inset). This high reactivity corresponds to the interface with uS19 observed in our cryo-EM structure, demonstrating that these nucleotides are likely flexible, and we hypothesise that this region may act as a hinge, allowing movements between dIVc_1_ and dIVc_2_ prior to ribosome binding. Interestingly, the pattern of SHAPE reactivity is very similar in the presence or absence of PCBP2 (**Figure 7B**), suggesting that PCBP2 does not induce large rearrangements at the level of secondary structure. It may therefore function in a more subtle way, perhaps by adjusting the tertiary positions of dIVc_1_, dIVc_2_ and dIVc_3_ for productive interaction with the ribosome (Beckham *et al*., 2020).

**Figure 7.**
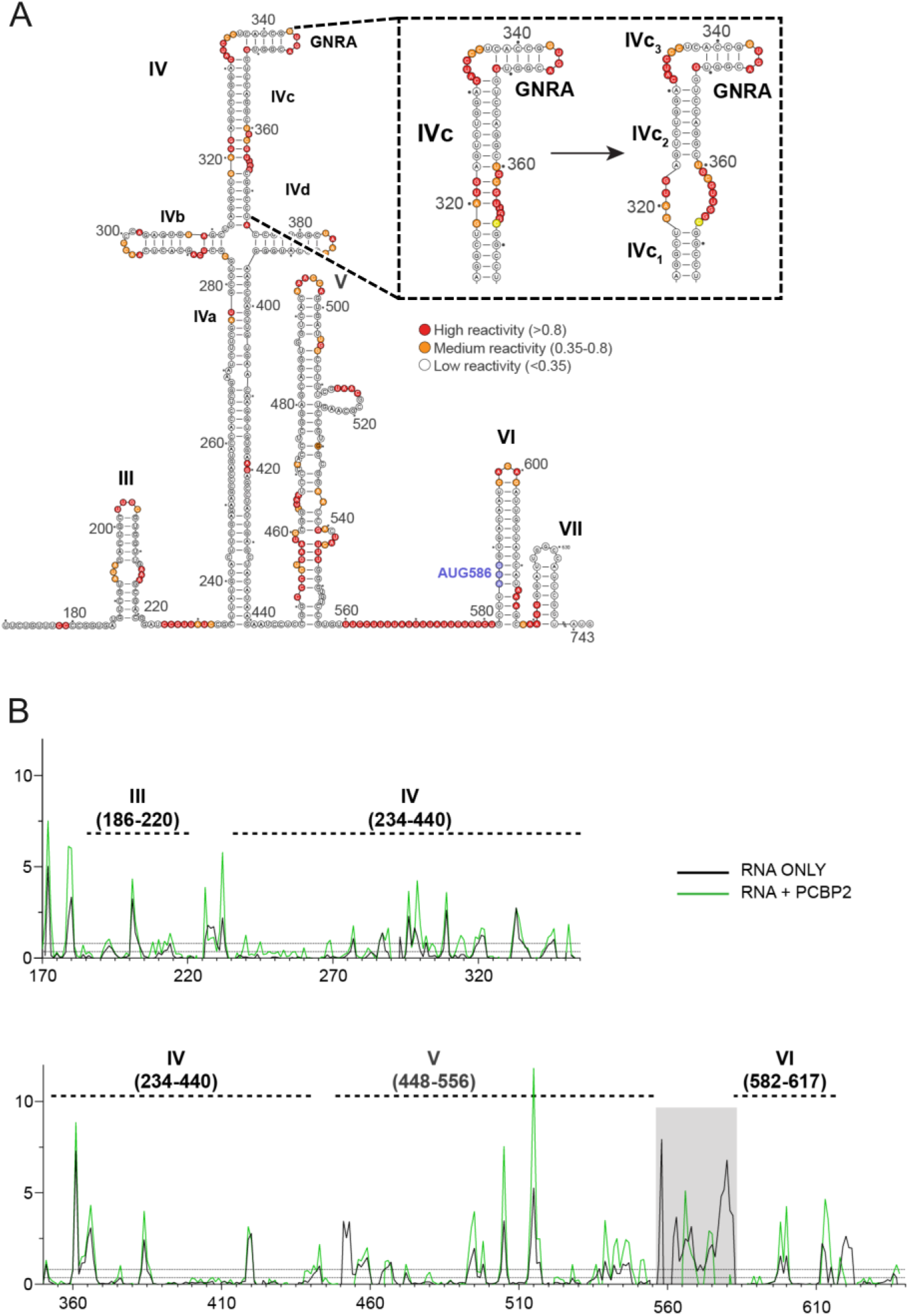
PCBP2 has limited influence on IRES secondary structure. **A,** SHAPE reactivities 0-0.35, 0.35-0.8 and >0.8 graded as low, medium and high, respectively, overlaid on the sequence of the PV IRES. **B,** Average SHAPE reactivity traces from 2-3 separate experiments in the presence or absence of PCBP2. The region in grey was variable between experiments.

### Disruption of dIVc contacts with the ribosome, tRNA or PCBP2 inhibit IRES activity and virus replication

To evaluate the significance of the dIVc contacts described above, we tested the effects of structure-guided mutations on initiation *in vitro*. First, we systematically removed domains from the 5′ end of the IRES (**Figure EV5A**) and assessed 48S complex formation at AUG_586_ by toeprinting (**Figure EV5B, Appendix Figure S1B**). Truncation of dI-III is tolerated, however removal of dI-IV or dI-V inhibits 48S assembly. In the latter case, we see multiple non-specific toeprints, possible due to off-pathway ribosome loading given that there are only eight nucleotides between AUG_586_ and the 5′ end. These results are consistent with previous literature: dI comprises the cloverleaf structure, involved in RNA replication rather than translation (Andino *et al*, 1990). Domain III can be removed entirely without affecting IRES activity in HeLa cells (Dildine & Semler, 1989). The role of dII is thought to be for binding to ITAFs UNR (Anderson *et al*, 2007) and possibly GARS (Andreev *et al*, 2022), but these are not required for initiation *in vitro* (Sweeney *et al*., 2014). However, removal of dIV is strongly inhibitory to 48S complex formation, consistent with our structural data.

To examine this further, we designed nine mutants to affect intra- and inter-molecular contacts observed in our structure (**Figure EV5C**). Mutants (1) and (2) are predicted to prevent GNRA interaction by shortening the C-rich loop (1) and stem (2) in dIVc_3_. Mutants (3) and (4) directly disrupt the GNRA loop. Mutants (5)-(7) are designed to disrupt uS13 interaction via co-mutating C-G residues in dIVc_2_ to G-C (5), U-A (6), or deleting them entirely (7). Finally, mutants (8) and (9) are designed to affect interaction with uS19 via G to A mutation (8) or deletion of a bulged, unpaired G residue (9). Efficiency of 48S complex formation at AUG_586_ was assessed by toeprinting and quantified by densitometry, using RT stops at the dV boundary to normalise for cDNA loading. Results are expressed relative to a wild-type control (**Figure EV5D,E and Appendix Figure S1C**). All mutations had a deleterious effect on 48S assembly *in vitro*. Disruption of the GNRA-tRNA interaction (mutants 1-4) had the greatest impact (∼20% wt), followed by targeting the uS19 contacts (mutants 8-9; ∼30-50% wt). Changing the identity of a base-pair in dIVc_2_ adjacent to the N105 interaction site in uS13 had a relatively mild effect (mutants 5-6, ∼65% wt), whereas removing this base-pair (mutant 7) reduced initiation to ∼40% wt levels.

We next investigated the effects of these mutations in viral IRES-dependent translation and replication in cells. To do this, we used a recently developed reverse genetics system for the closely related enterovirus CVA13 (O’Connor *et al*, 2026), which belongs to the same species as PV (*Enterovirus coxsackiepol*). The high sequence conservation between these two strains (87% pairwise identity for dIVc) makes CVA13 a suitable surrogate for PV (**Figure 8A**) and we engineered equivalent mutations to those described above in a CVA13 background (**Figure 8B,C**).

**Figure 8.**
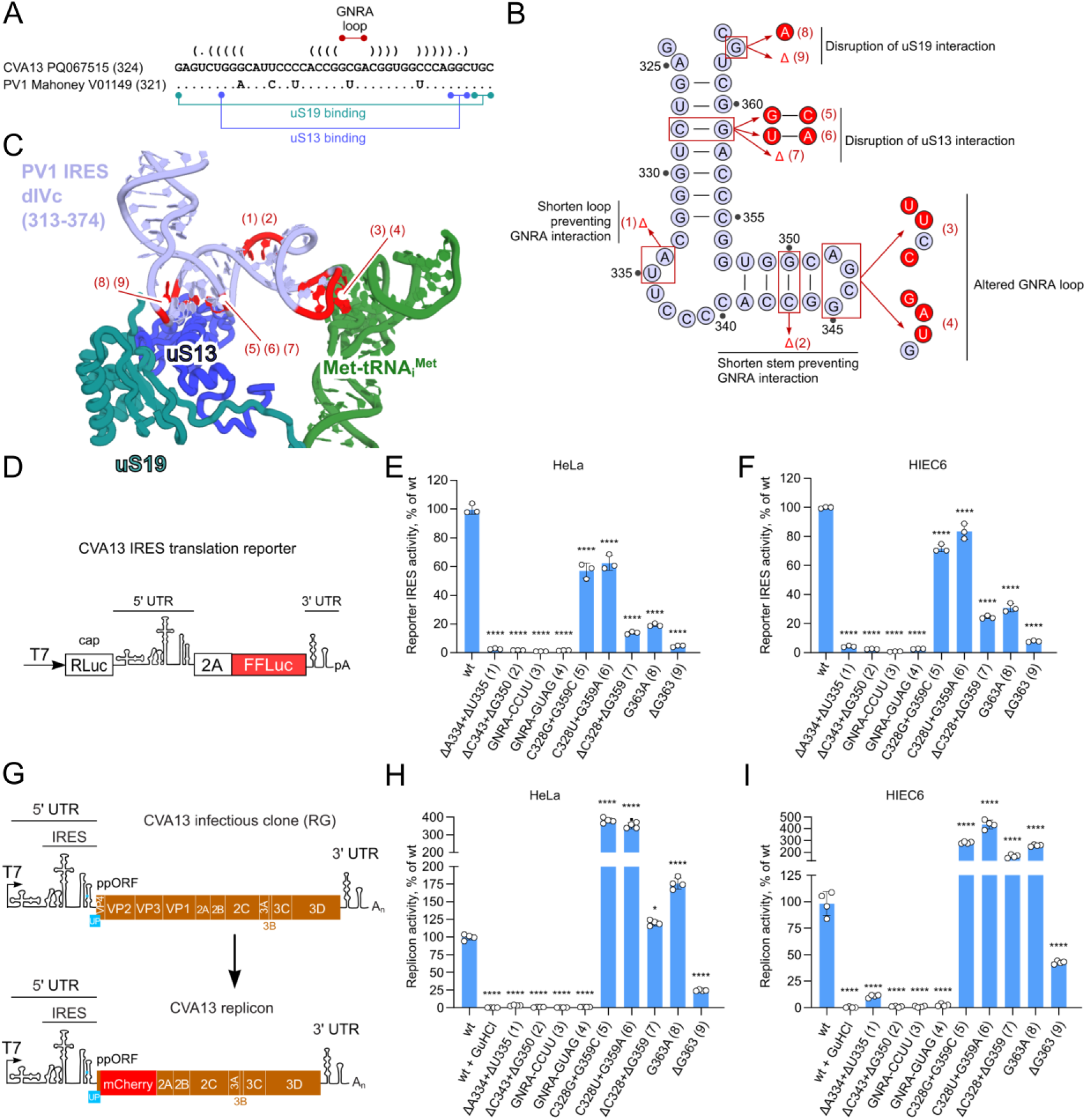
dIVc mutations in CVA13 have a severe impact on virus translation and replication. **A,** IRES domain IVc RNA sequence comparison between coxsackievirus A13 (CVA13) and poliovirus 1 (PV1) Mahoney strain (both are Enterovirus coxsackiepol species). **B,** Location of mutations on the PV dIVc structure are indicated (red). **C,** Location of mutations on the equivalent CVA13 dIVc sequence are indicated (red). **D,** Schematic representation of the dual luciferase reporter used to measure IRES-dependent (firefly luciferase) relative to cap-dependent (Renilla luciferase) translation (O’Connor et al., 2026). **E-F,** Analysis of IRES relative to cap-dependent activity for wt (100%) and indicated mutant reporters measured in (E) HeLa and (F) HIEC6 cells (n=3 biological replicates). P = 6×10-15, 6×10-15, 6×10-15, 6×10-15, 1×10-13, 2×10-12, 6×10-15, 6×10-15, 6×10-15 (panel E, left to right) and P = 6×10-15, 6×10-15, 6×10-15, 6×10-15, 3×10-12, 6×10-8, 6×10-15, 6×10-15, 6×10-15 (panel F, left to right) using one-way ANOVA with Dunnett’s post-hoc test against wt control. **G,** Schematic representation of the CVA13 replicon derived from the reverse genetics-based plasmid (O’Connor et al., 2026). The structural proteins are replaced by mCherry while preserving cleavage sites. **H-I,** Analysis of activities of wt (100%) and indicated mutant replicons tested in (H) HeLa and (I) HIEC6 cells (n=4 biological replicates). Guanidine hydrochloride (GuHCl) is included as a well-established inhibitor of the viral helicase that prevents replication (wt + GuHCl). P is <10-15, <10-15, <10-15, <10-15, <10-15, <10-15, <10-15, 2×10-2, 9×10-14, 9×10-14 (panel H, left to right) and 7×10-10, 1×10-8, 8×10-10, 8×10-10, 1×10-9, <10-15, <10-15, 6×10-6, <10-15, 8×10-5 (panel I, left to right) using one-way ANOVA with Dunnett’s post-hoc test against wt control. Data (E,F,H,I) are mean ± SD. **** p < 0.001, * p < 0.05; ns, nonsignificant.

First, we evaluated IRES-dependent translation using the recently developed CVA13-based IRES reporter system, in which firefly luciferase activity reports on ppORF translation (**Figure 8D**) (O’Connor *et al*., 2026). We performed experiments in both HeLa cells and human intestinal epithelial cells (HIEC6) as a more relevant enteric virus model system. As shown in **Figure 8E,F**, all mutations affecting dIVc_3_ including the GNRA loop itself (mutants 3-4) or its context (mutants 1-2) were highly detrimental to translation (<5% in both cell lines). Disruption of the uS13 contact (mutants 5-6) reduced translation by ∼20% while shortening dIVc_2_ reduced translation to ∼25% of wt (mutant 7). A similar pattern was observed for uS19 contact: the deletion of G resulted in <8% IRES-dependent translation. Taken together, the GNRA loop has the strongest impact on the IRES-dependent translation, followed by important interactions with uS19 and uS13. These results are in good agreement with our *in vitro* data.

Next, we assessed the impact of these mutations in a replication-competent system. Following viral entry, IRES-dependent translation is the crucial first step in the virus life cycle, followed by RNA replication performed by newly translated non-structural proteins and recruited host factors (proteins, membranes). To enable comparison with the IRES activity assays described above, we developed a CVA13 replicon system comprising an intact CVA13 genome with the structural proteins replaced by an mCherry coding sequence (**Figure 8G**). This replicon provides a direct measurement of CVA13 replication but lacks a packaging step. Following the translational assay scheme (**Figure 8E,F**), we performed replicon assays in HeLa and HIEC6 cell lines (**Figure 8H,I**). Consistent with translation assay results, all four GNRA mutants (1-4) resulted in complete suppression of replication, comparable to levels of replication-deficient control (wt + GuHCl). This confirms the ultimate importance of the GNRA sequence and its position for efficient translation, which is a prerequisite for RNA replication. Interestingly, mutants 5-8 resulted in increased replication levels in HeLa and HIEC6 cells. This can be explained by competition between translation and replication processes over the same RNA molecule (Gamarnik & Andino, 1998). In the scenario of a milder translational defect and reduced ribosome load, the RNA is more available to act as a template of replication, resulting in overall higher genome replication efficiency (**Figures 8E,F,H,I**). Consistently, the deletion mutant affecting uS13 interaction (mutant 7) results in lower replication efficiency when compared to mutants 5 and 6 in both cell lines, following the translation assay pattern, but still outperforms wt replication. The deletion mutant affecting uS19 interaction (mutant 9) had a more severe translation defect and subsequently more severe replication defect, confirming the importance of this host-pathogen interaction in both tested systems (**Figures 8H,I**).

Finally, we used our CVA13 reporter and replicon systems to test the importance of the putative interface between the C-rich loop in dIVc_3_ and PCBP2 KH1 (**Figure 6C,D**). We made a series of C-U substitutions in this region (CCCC-CCUC, CCCC-CUUC and CCCC-UUUC, **Figure 9A**), all of which were designed to preserve the structure of the IRES but increasingly inhibit PCBP2 binding. Consistent with this, we observed progressive defects in IRES-dependent translation in both HeLa and HIEC6 cells, with the severity of the effect linked to the number of cytosine substitutions (**Figure 9B**). The most severe mutation (UUUC) reduced translation to <5% wt levels and also suppressed replication to ∼20% and ∼45% wt in HeLa and HIEC6 cells, respectively (**Figure 9C**). Mutations with a milder impact on translation (CCUC and CUUC) were seen to increase genome replication, as explained above (**Figure 9B,C**).

**Figure 9.**
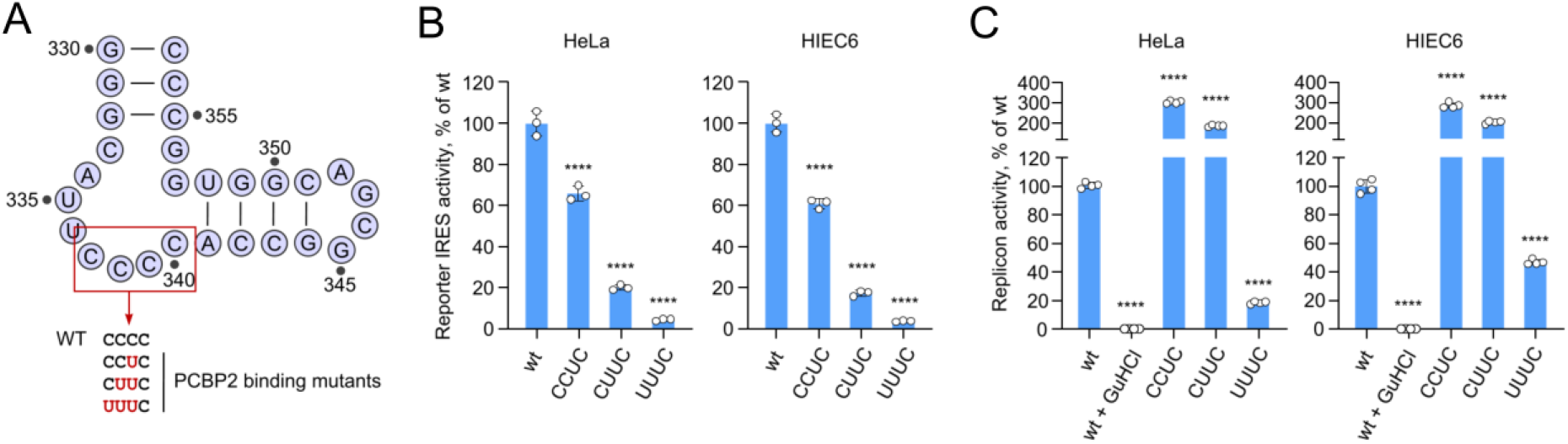
PCBP2-binding mutations in dIVc3 affect CVA13 translation and replication. **A**, IRES domain IVc3 RNA sequence with indicated mutations designed to disrupt PCBP2 binding. **B**, Analysis of IRES relative to cap-dependent activity for wt (100%) and indicated mutant reporters measured in (left) HeLa and (right) HIEC6 cells (n=3 biological replicates). P = 6×10-6, 9×10-9, 2×10-9 (left to right, HeLa) and 2×10-7, 6×10-10, 2×10-10 (left to right, HIEC6) using one-way ANOVA with Dunnett’s post-hoc test against wt control. **C**, Analysis of activities of wt (100%) and indicated mutant replicons tested in (left) HeLa and (right) HIEC6 cells (n=4 biological replicates). P = 10-15, <10-15, 3×10-14, 3×10-14 (left to right, HeLa) and 5×10-12, <10-15, 5×10-12, 4×10-8 (left to right, HIEC6) using one-way ANOVA with Dunnett’s post-hoc test against wt control. Data (B,C) are mean ± SD. **** p < 0.001.

Taken together, we validated our structural findings three ways: first *in vitro*, and secondly in cells using two independent systems that are directly affected by impaired translation of virus RNA. The results are consistent, confirming that efficient IRES-dependent translation is a prerequisite for successful RNA replication. Moreover, the relative translational output of different CVA13 mutants in cells (**Figure 8E,F**) matches the relative efficiency of 48S assembly on PV-AUG_586_-good mutants *in vitro* (**Figure EV5D,E**) suggesting that the observed translational defects in cells can be explained by impaired initiation. The most significant role can be assigned to the GNRA sequence, followed by interfaces with uS19, then uS13 (**Figure 8**). We also confirm the importance of the dIVc_3_-PCBP2 interaction in IRES-dependent translation and subsequent replication.

### Enterovirus dIVc is likely to be structurally conserved

We next explored to what extent the key IRES sequences identified here are conserved amongst enteroviruses. To do this, we aligned dIVc regions from multiple enteroviruses, including wild-type polioviruses (PV1, PV2, PV3), Sabin vaccine strains, CVA13 Flores strain (*Enterovirus coxsackiepol* species), three common *Enterovirus alphacoxsackie* strains (EV-A71, CVA6, CVA16), and three prevalent *Enterovirus betacoxsackie* strains (E30, E11, CVB6) (**Figure 10**). Despite the overall low primary sequence conservation across these 14 enterovirus strains, the overall secondary structure of dIVc was consistently preserved. Nearly all variations in this region were compensated by covariant substitutions that restored base pairing at key dIVc contacts. Furthermore, a C-U variation at the start of the C-rich loop in dIVc_3_ is often accompanied by a U-C change three nucleotides downstream, thereby maintaining at least three consecutive cytosines for PCBP2 binding. These findings support the existence of evolutionary conserved RNA structure that underlies Type 1 IRES function across diverse enteroviruses species.

**Figure 10.**
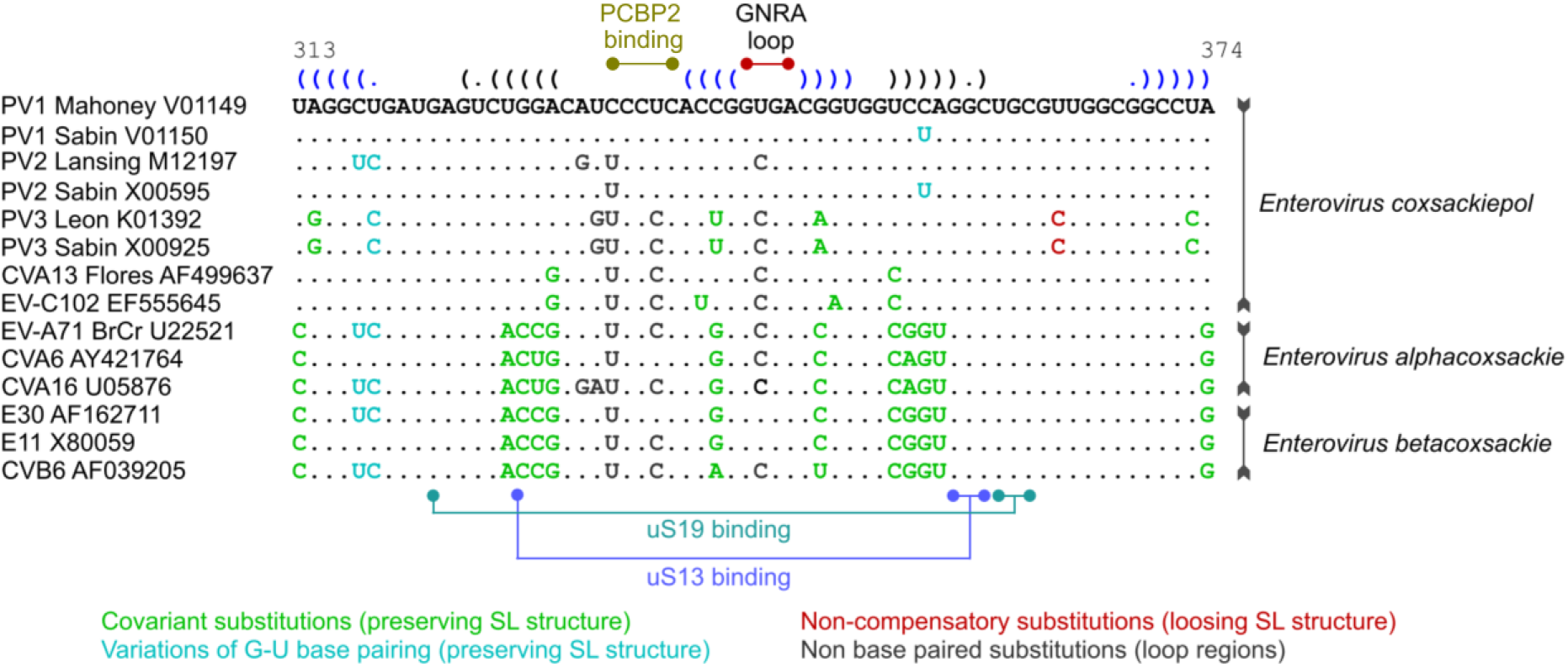
Alignment and structural conservation of enterovirus IRES dIVc. Sequence alignments for Enterovirus coxsackiepol (PV1, PV2, PV3, CVA13, EV-C102), alphacoxsackie (EV-A71, CVA6, CVA16) and betacoxsackie (E30, E11, CVB6) species. Substitutions that preserve SL structures are shown green (C-G/A-U) or blue (G-U variations). Substitutions that break SL structure are shown in red. Non base paired substitutions are indicated in grey.

Beyond enteroviruses, recent structural studies of 48S initiation complexes on the Type 2 EMCV IRES (Bhattacharjee *et al*, 2026; Data ref: PDB 8SUP, 2026; Das & Hussain, 2026 Data ref: PDB 9UZK, 2026; Data ref: PDB 9UZL, 2026) have revealed unexpected similarity with the PV Type 1 IRES, implying that some aspects of initiation may be conserved amongst divergent picornaviruses (**Appendix Figure S8**). Cryo-EM structures of closed 48S complexes at EMCV AUG_834_ show density for nucleotides 518-600 of domain I (equivalent to PV dIV) binding to the translation machinery in a similar way. EMCV Ih2 interacts with uS13 in a manner analogous to PV dIVc_2_, whilst the GNRA motif (EMCV GCGA 547-550, located at the tip of Ih3), contacts the TѰC loop of the tRNA_i_. However, contributions from EMCV Ih4 (absent in PV) extend the interface with the 40S head, additionally involving uS13 residues E112, K116, R118 and uS19 residues N103, Q104, I121 and P125. As a result, the buried surface area is larger in EMCV (∼1125 Å^2^) than PV (∼587 Å^2^) (**Appendix Figure S8**), perhaps contributing to a more stable 40S-IRES interaction (Chamond *et al*, 2014).

## Discussion

### An intermolecular RNA tertiary interaction underpins enterovirus translation

IRESs are essential for replication of numerous pathogenic human and livestock viruses, enabling continued viral protein synthesis in the face of depleted global host cell translation. Until now, structural information on the organisation of the high complexity Type 1 IRES on the ribosome has remained elusive. Here, we present the first structure of an early 48S initiation complex on the PV Type 1 IRES and reveal a critical intermolecular contact between the highly conserved GNRA loop at the tip of dIVc, and a helix on the initiator tRNA. Intramolecular GNRA interactions with minor grooves are common tertiary structural motifs that drive the folding of complex RNAs (Bottaro & Lindorff-Larsen, 2017; Fiore & Nesbitt, 2013) including group I and II introns (Adams *et al*, 2004; Costa *et al*, 1997; Costa & Michel, 1995; Michel & Westhof, 1990), RNAse P (Massire *et al*, 1997; Tanner & Cech, 1995) and ribosomal RNA (Ban *et al*, 2000; Belanger *et al*, 2004; Wimberly *et al*, 2000). However, fewer examples exist of heterotypic intermolecular GNRA-minor groove interactions, and we can now extend this list to include virus-host interactions. Our translation and replication assays show that deviation from the optimal GNRA sequence completely abolishes IRES activity and inhibits virus replication, consistent with previous observations. Notably, the effects of mutating uS13 and uS19 interaction surfaces are less severe, implicating the GNRA-tRNA_i_ interface as the most biologically significant.

The GNRA loop is highly conserved, and essential for the activity of many Type 1 and Type 2 IRESs (Fernández-Miragall & Martínez-Salas, 2003; Fernández *et al*., 2013; Fernández *et al*., 2011; López de Quinto & Martínez-Salas, 1997; Nateri *et al*., 2000; Robertson *et al*., 1999). Both share a common structural arrangement: a large domain bearing an apical GNRA loop precedes an eIF4G/eIF4A binding stem, followed by a polypyrimidine rich sequence preceding an initiation codon. However, whilst initiation may proceed via a conceptually similar mechanism, their sequences are unrelated and important virus-specific differences exist. For example, Type 1 IRESs require binding of eIF4G and eIF3 to promote translation initiation, an interaction dispensable for Type 2 IRES function (Sweeney *et al*., 2014).

The relationships between the GNRA motif, IRES conformation and ITAF requirements are complex. In the divergent cadicivirus Type 1 IRES, an intact GNRA sequence is strictly required for IRES activity, and for the ability of PCBP2 to act as an ITAF (Asnani *et al*., 2016). Structure probing of this IRES showed no major structural differences between wild-type or GNRA-mutated RNAs. In contrast, mutation of the GNRA loop in the Type 2 FMDV IRES led to a reorganisation of the I-domain (equivalent to PV dIV) (Fernández-Miragall & Martínez-Salas, 2003; Fernández-Miragall *et al*, 2006), suggesting that the GNRA loop may interact with an alternative component of the IRES. This may be related to the need for different ITAFs: EMCV and FMDV Type 2 IRESs do not use PCBP2, instead requiring PTB and ITAF45 (Bellucci *et al*, 2025; Hellen *et al*, 1993) both of which have been reported to induce structural changes in their cognate IRESs (Yu *et al*, 2011a). Indeed, while the specific site(s) of ITAF45 interaction have not been determined, multiple unique sites of PTB interaction with the EMCV IRES were described (Kafasla *et al*, 2009), consistent with a greater impact on IRES structure. In contrast, PCBP2 binding is localised to the apex of dIV in the Type 1 IRES (Sweeney *et al*., 2014). The Type 5 aichivirus-like IRESs exhibit further differences still, comprising a hybrid Type 1 IRES dIV-like structure and Type 2 IRES J/K domain-like structure. In contrast to both Type 1 and 2 IRESs, deviation from the GNRA tetraloop in the Type 5 IRES in some cases led to an increase in IRES activity (Yu *et al*, 2011b). While the location of the initiation codon in a stable secondary structure explains the stimulatory role for the translation initiation factor DHX29, how PTB promotes Type 5 IRES activity remains to be determined.

### PCBP2 likely primes dIVc for productive ribosome encounter

PCBP2 regulates gene expression at transcriptional and posttranscriptional levels and is increasingly linked to cancer progression (Yuan *et al*, 2021). The tripartite KH domain architecture confers a high degree of modularity in RNA binding: specificity for C-rich sequences, combined with a long, unstructured linker between KH1/KH2 and KH3 domains provides a mechanism to stabilise tertiary RNA structure by binding to distal RNA elements. *In vitro*, PCBP2 binds to dIV and is necessary for formation of 48S complexes on several Type 1 IRESs, including PV, enterovirus 71 and bovine enterovirus (Beckham *et al*., 2020; Sweeney *et al*., 2014). Interestingly however, we did not observe density for PCBP2 in our structures, suggesting that during AUG recognition, PCBP2 is no longer required to maintain the network of interactions between dIVc, uS13, uS19 and tRNA_i_ after they have formed. Whilst PCBP2 binding to dIVc_3_ is essential for efficient translation (**Figure 9**), our SHAPE data revealed no major differences to the IRES secondary structure in the presence or absence of PCBP2 (**Figure 7**). A caveat of our SHAPE analysis is that it represents an ensemble of molecules at equilibrium during a reaction window of 50 minutes. Although PCBP2 was present in molar excess, depending on its on- and off-rates, and the nature of the interactions, very rapid changes may not be detectable. We also note that the region of differential SHAPE reactivity observed between dV and dVI was not consistent between replicates, and therefore cannot be ascribed to PCBP2 binding. Nevertheless, taken together, our data implies that PCBP2 may function at an earlier initiation step than visualised in our structures, perhaps by altering the relative positioning of dIVb, dIVd and dIVc for efficient GNRA presentation.

We and others have also shown an interaction between GARS (Andreev *et al*, 2012; Sweeney *et al*., 2014) and La (Meerovitch *et al*., 1993) proteins with the PV IRES, although unlike PCBP2, these did not promote 48S initiation complex assembly in an *in vitro* reconstituted system. La did however alter the structure of the viral RNA in the domain V-VI region in the presence and absence of PCBP2 (Sweeney *et al*., 2014) consistent with a potential role in refining IRES activity. Whether other Type 1-like IRESs that use PCBP2 as an ITAF control genome usage in a similar manner remains to be determined.

### Translation-replication dynamics in enterovirus life cycle

In the picornavirus life cycle, translation and replication compete for access to the same RNA molecule (Gamarnik & Andino, 1998). This competition is essential for efficient virus replication and is regulated by a combination of cellular and viral factors. Specifically, the viral 3CD/3C protease plays a key role in this process by cleaving PCBP2, inhibiting the activity of the GNRA loop and shifting the balance in favour of RNA replication over translation (Holmes & Semler, 2019; Perera *et al*, 2007). This may have evolved as a mechanism for temporal control of genome usage, optimising replication efficiency (Chase *et al*, 2014).

Translation-replication dynamics are tightly controlled in each infected cell and ultimately determine the outcome of infection (Boersma *et al*, 2020). Consequently, defects in translation can profoundly disrupt RNA replication. Severe translational impairments can completely suppress replication, as observed for GNRA mutants (**Figure 8**). In contrast, mild defects in translation, such as those observed for uS13 binding mutants, may still allow efficient amplification of RNA genome, as multiple rounds of RNA replication compensate for the reduced translation. However, this imbalance is generally detrimental to the full viral life cycle, since productive infection requires sufficient translation to generate the structural proteins necessary for RNA packaging and virion assembly, steps that are absent in the replicon system.

### Towards a multi-step model for initiation on the Type 1 IRES

Our structures suggest that the mechanism by which ribosomes initiate on Type 1 IRESs is closer to canonical cap-dependent initiation than to mechanisms of smaller IRESs for which structures are available. CrPV-like and HalV-like IRESs directly bind to specific sites in the A- and P-sites of the 40S subunit, enabling elongation-competent ribosomes to assemble at a fixed position on the viral mRNA without an initiator tRNA (Abaeva *et al*., 2020; Acosta-Reyes *et al*., 2019; Fernández *et al*., 2014; Muhs *et al*., 2015; Murray *et al*., 2016; Pisareva *et al*., 2018; Spahn *et al*., 2004). Similarly, HCV-like IRESs form a network of interactions across the 40S surface, displacing eIF3 to position the initiator tRNA directly in the P-site, without scanning (Brown *et al*., 2022; Hashem *et al*., 2013; Quade *et al*., 2015; Yamamoto *et al*., 2015; Yamamoto *et al*., 2014; Yokoyama *et al*., 2019). In contrast, our observation of both ‘open’ and ‘closed’ 48S complexes during PV initiation is consistent with previous biochemical demonstrations of scanning and the requirement for eIF4A (Pause *et al*, 1994; Sweeney *et al*., 2014).

This implies that PV initiation likely proceeds in a regulated, step-wise manner through 43S recruitment, local scanning, AUG recognition and large subunit joining, although exactly where 43S complexes are internally loaded is not known. Interactions previously identified as essential for PV 48S complex formation, such as eIF4G-eIF3 (Sweeney *et al*., 2014) and dV-eIF4G-4A (de Breyne *et al*., 2009) may therefore only be needed for specific events. This could explain the lack of density for dV, eIF4G or eIF4A in our structures: instead, these elements may be required at an earlier step. Notably, in recent structures of 48S complexes on the EMCV Type 2 IRES, density for the J-K-St domain (equivalent to Type 1 dV), eIF4G or eIF4A was also not observed (Bhattacharjee *et al*., 2026; Das & Hussain, 2026), although structures of eIF4G-4A bound to the J-K-St domain in isolation reveal a preorganised RNA platform that binds directly to eIF4G independently of 40S or other eIFs (Imai *et al*, 2016; Imai *et al*, 2023). In PV, the idea of dV-4G-4A acting as a driving force for 43S recruitment is consistent with the absence of IRES density in our 40S classes and suggests that the dIVc-uS13-uS19 interface alone is insufficient to drive initiation. In a similar way, we note a minimal involvement of eIF3 in our ‘closed’ 48S complexes at AUG_586_. No contacts are observed between dIVc and any eIF3 subunit, and the dIVc-uS13-uS19-tRNA_i_ interactions can persist following the dissociation of eIF3.

What happens next? The dIVc interactions observed in our closed 48S complexes are incompatible with motions of tRNA following GTP hydrolysis by eIF2 (Petrychenko *et al*., 2025), the binding of eIF5B (Lapointe *et al*., 2022) or 60S joining (Holm *et al*., 2023). It will be intriguing to observe how these contacts are remodelled or displaced at later stages of 48S assembly, post GTP hydrolysis. Interestingly, eIF5B is cleaved by the 3C protease during enterovirus infection, separating the N-terminal domain from its GTPase and C-terminal domains (de Breyne *et al*, 2008) but it remains to be investigated how this affects IRES function.

In conclusion, our work describes a new molecular interaction that underpins enterovirus genome translation and is likely a general feature of all Type 1 IRESs. Very recent work also suggests that similar interactions drive initiation on the Type 2 EMCV IRES (Bhattacharjee *et al*., 2026; Das & Hussain, 2026), revealing a surprising degree of mechanistic similarity between divergent picornaviruses. Together these studies provide structural frameworks to rationalise decades of previous virological and biochemical work, highlighting the untapped potential of RNA-RNA tertiary interactions as targets for novel antivirals.

## Methods

### Methods and Protocols

#### Recombinant protein expression and purification

Protein-coding sequences were cloned into either pET-28 (eIF1, eIF4Gm, PCBP2, aminoacyl tRNA synthetase, eIF3j) or pET-15b (eIF4A, eIF4B) vectors, with an N-terminal His_6_ tag for immobilised metal ion affinity chromatography (IMAC). Briefly, plasmids were transformed into *E.coli* BL21 cells and grown shaking (37°C, 210 rpm) in Luria-Bertani (LB) medium supplemented with 50 μg/mL kanamycin or 100 μg/mL ampicillin, as appropriate. Expression was induced at an optical density (600 nm) of 0.5 by the addition of 1mM IPTG (3 h, 37°C, 210 rpm) prior to pelleting by centrifugation (4000 x g, 4°C, 10 min) and storage (-20°C).

Cell pellets were thawed and resuspended in ice-cold lysis buffer (20 mM Tris-HCl pH 7.5, 300 mM KCl, 10% v/v glycerol supplemented with 1.0 mM AEBSF, 0.7 µg/mL Pepstatin A, 0.5 µg/mL Leupeptin and 25 µg/mL DNAse I prior to lysis in a cell disruptor at 24 kPSI (Constant Systems). Lysates were clarified (39,000 x g, 4°C, 30 min) and incubated in a gravity flow column (BioRad) with 800 μL Ni-NTA Sepharose 6 Fast Flow beads (Cytiva) pre-equilibrated in buffer (20 mM Tris-HCl pH 7.5, 300 mM KCl, 20 mM imidazole, 10% v/v glycerol). Beads were washed with 10 mL 20 mM Tris-HCl pH 7.5, 800 mM KCl, 20 mM imidazole, 10% v/v glycerol, followed by 20 mL of 20 mM Tris-HCl pH 7.5, 100 mM KCl, 20 mM imidazole, 10% v/v glycerol. Proteins were eluted by addition of 20 mM Tris-HCl pH 7.5, 100 mM KCl, 300 mM imidazole, 10% v/v glycerol, and dialysed (10K MWCO, 4°C, 16h) to remove imidazole. Samples were then applied to either a MonoQ 5/50 column or MonoS 5/50 column (Cytiva) and eluted by a linear salt gradient from 100 mM KCl to 500 mM KCl. Fractions containing proteins of interest were identified by SDS-PAGE (**Figure EV1**), pooled, exchanged into storage buffer (20 mM Tris-HCl pH 7.5, 100 mM KCl, 0.1mM EDTA, 10% glycerol) and concentrated to at least 1.0 mg/mL using a 30K Amicon Ultra-15 concentrator (Millipore) prior to snap-freezing in liquid nitrogen and storage at -80 °C.

#### Purification of human ribosome subunits

Purification of human 40S and 60S ribosomal subunits was performed based on published methods (Fraser *et al*., 2007; Pisarev *et al*, 2007) with modifications. HeLa cytoplasmic extract (Ipracell, 168 mL) in 10 mM HEPES-KOH pH 7.9, 0.5 mM Mg(OAc)_2_, 10 mM KOAc, 0.5 mM DTT (Ipracell) was thawed and supplemented with 0.5 mg/mL Pefabloc SC (Roche), 0.7 µg/mL pepstatin A, 0.5 µg/mL leupeptin and 3.5 mM Mg(OAc)_2_, prior to clarification (20,000 x g, 4°C, 20 min) and layering onto dual 0.5 M and 0.7 M sucrose cushions in 20 mM Tris-HCl pH 7.5, 4.0 mM MgCl_2_, 50 mM KCl, 2.0 mM DTT. Polysomes were pelleted by ultracentrifugation in a Ti-45 rotor (234,000 x g, 4°C, 4h), rinsed with ice-cold water and resuspended in buffer A (20 mM Tris-HCl pH 7.5, 4.0 mM MgCl_2_,50 mM KCl, 2.0 mM DTT). To split polysomes, puromycin was added to a final concentration of 1.5 mM, incubated (4°C for 10 min, then 37°C for 10 min) and stirred on ice (30 min). 4.0 M KCl was slowly added dropwise to a final concentration of 0.5 M KCl prior to ultracentrifugation in a Ti-45 rotor (234,000 x g 4°C, 4.5 h). The “ribosomal salt wash” (RSW) supernatant was kept for purification of eIF3 and eIF2. Ribosome pellets were washed with ice-cold water and resuspended in buffer B (20 mM Tris-HCl pH 7.5, 2.0 mM DTT, 4.0 mM MgCl_2_, 0.5 M KCl) before centrifugation through a 10-30% (w/v) sucrose gradient in buffer B (140,000 x g, 4°C, 16h) to separate 40S and 60S subunits. Fractions containing uniquely 40S or 60S subunits were identified by SDS-PAGE (**Figure EV1**), pooled and exchanged into buffer C (20 mM Tris-HCl pH 7.5, 2.0 mM DTT, 4.0 mM MgCl_2_, 100 mM KCl, 0.25 M sucrose) using a 100K MWCO Amicon Ultra-15 concentrators (Millipore). Subunits were snap-frozen in liquid nitrogen at concentrations of A_260_ 25-60 U/mL (40S) or A_260_ 60-120 U/mL (60S) and stored at -80 °C.

#### Purification of human eIF3

Purification of human eIF3 was based on published methods (Pisarev *et al*., 2007) with modifications. Finely powdered (NH_4_)_2_SO_4_ was slowly added to the RSW fraction (stirring on ice, 30 min) to generate a 0-40% fraction (242 g/L RSW). This was pelleted by centrifugation (50,000 x g, 4°C, 20 min) and dissolved in 5.0 mL buffer D (20 mM Tris-HCl pH 7.5, 100mM KCl, 2.0 mM DTT, 0.1 mM EDTA, 10% w/v glycerol) and dialyzed against 1.0 L buffer D (10K MWCO, 4°C, 16h). This fraction was then applied to an 11 mL DEAE Sepharose anion exchange column (Cytiva) pre-equilibrated in buffer D. A step to 250 mM KCl over five column volumes (CV) was used to elute eIF3, eIF4F, eIF4B and small amounts of eIF2. Eluate was adjusted to a KCl concentration of 100 mM before application to a Capto HiRes S 10/100 column (Cytiva) equilibrated in buffer D. A linear KCl gradient from 100 mM to 1.0 M (30 CV) was used to elute eIF3-containing fractions, which were pooled and layered onto a 10-30% (w/v) sucrose gradient in buffer E (20 mM Tris-HCl pH 7.5, 400 mM KCl, 2.0 mM DTT, 0.1 mM EDTA, 10% w/v glycerol). Ultracentrifugation in an SW-41 rotor (287,000 x g, 4°C, 22 h) was performed to separate eIF3 from eIF4G, as determined by SDS-PAGE and Western blotting of fractions with rabbit α-human eIF3D (Proteintech, 10219-1-AP), rabbit α-human eIF4G (Cell Signalling Technology, C45A4) and rabbit α-human eIF4E (Cell Signalling Technology, C46H6) antibodies. Suitable fractions were exchanged into buffer D and applied to a MonoQ 5/50 anion exchanger (Cytiva) equilibrated in buffer D. A linear gradient from 100 mM → 500 mM KCl (20 CV) was used to elute eIF3. Fractions containing eIF3 were pooled and exchanged into buffer D using a 100K MWCO Amicon Ultra-15 concentrator (Millipore) prior to snap-freezing in liquid nitrogen and storage at -80 °C (**Figure EV1**).

#### Purification of human eIF2

Purification of human eIF2 was based on published methods (Pisarev *et al*., 2007) with modifications. After pelleting the 0-40% (NH_4_)_2_SO_4_ fraction, further (NH_4_)_2_SO_4_ was added to the supernatant (62 g/L RSW) to generate a 40-50% fraction. This was pelleted, exchanged into buffer D, and subjected to DEAE Sepharose and Capto HiRes S 10/100 chromatography as described above. Fractions containing eIF2 were pooled and adjusted to 100 mM KCl, before being applied to a MonoQ 5/50 anion exchanger (Cytiva) equilibrated in buffer D. A linear gradient (20 CV) from 100 mM → 500 mM KCl was used to separate eIF2 from eIF5B. eIF2-containing fractions were then exchanged into buffer F (20 mM HEPES-KOH pH 7.5, 100 mM KCl, 100 mM K_2_HPO_4_/KH_2_PO_4_, 1.0 mM DTT, 10% v/v glycerol) and applied to a CHT5-1 ceramic hydroxyapatite column (BioRad) that had been pre-equilibrated in the same buffer. eIF2 was eluted using a linear gradient from 100 → 400 mM K_2_HPO_4_/KH_2_PO_4_ (5 CV). Fractions containing pure eIF2 were pooled and exchanged into buffer D using a 30K MWCO Amicon Ultra-15 concentrator (Millipore) before snap-freezing in liquid nitrogen and storage at -80 °C (**Figure EV1**).

#### RNA transcription for reconstitution

Sequences encoding the PV-AUG_586_-good IRES were cloned into pUC57 by PCR amplification, restriction digest and ligation. RNA was produced by run-off *in vitro* transcription using 10 µg of NdeI-linearised pUC57 template and T7 RNA polymerase (prepared in-house, 37°C, 4h). Reactions comprised 40 mM HEPES-KOH pH 7.5, 32 mM Mg(OAc)_2_, 40 mM DTT, 2.0 mM spermidine, 5.0 mM ribonucleoside triphosphates (rATP, rGTP, rCTP, rUTP). Template DNA was removed by digestion with RNAse-free DNAse I (Thermo, 37°C, 1h) prior to extraction in acid phenol:chloroform, ethanol precipitation and reconstitution in distilled water. To remove unincorporated nucleotides RNA was exchanged into 20 mM Tris-HCl pH 7.0, 1.0 mM EDTA using a 5.0 mL HiTrap Desalting column (Cytiva). Integrity of purified RNAs was confirmed by Urea-PAGE, and RNAs were stored at -80 °C.

#### Charging of initiator tRNA with methionine

Calf liver tRNA (Promega) was charged with L-Methionine using aminoacyl tRNA synthetase (aa-tRS, prepared in-house, 37°C, 1.5 h). Reactions of 200 μL comprised 2.0 mg total tRNA, 60 μg aa-tRS, 40 mM Tris-HCl pH 7.5, 15 mM MgCl_2_, 4.0 mM ATP, 1.0 mM CTP, 75 μM L-methionine. Charged tRNAs were extracted in acid phenol:chloroform, applied to a G-50 Illustra spin column (GE), ethanol precipitated and reconstituted in 20 mM Tris-HCl pH 7.0, 1.0 mM EDTA.

#### Termination of primer extension assays

<u>T</u>ermination <u>o</u>f primer <u>e</u>xtension assays (toeprinting) were performed as described (Pisarev *et al*., 2007). Briefly, a DNA primer was labelled with [γ-32P] dATP (6000 Ci/mmol; Revvity) using T4 polynucleotide kinase (3′ phosphatase minus, NEB, 37°C, 20 min). Unincorporated radiolabel was removed using G-50 Illustra spin columns (GE) according to the manufacturer’s instructions. Separately, complexes were assembled by mixing 2.0 pmol 40S, 20 pmol PCBP2, 5.0 pmol eIF4Gm, 5.0 pmol eIF4B, 10 pmol eIF4A, 10 pmol eIF1A, 1.0 pmol eIF3, 2.0 pmol eIF2, 200 pmol tRNA in buffer (final concentrations 20 mM Tris-HCl pH 7.5, 60 mM KCl, 2.5 mM MgCl_2_, 2.0 mM DTT, 0.25 mM spermidine, 0.5 mM GTP, 2.0 mM ATP). Complexes were equilibrated (37°C, 5 min) prior to addition of 2.0 pmol PV-AUG_586_-good IRES RNA and further incubation (37°C, 10 min). For primer extension, 20 μL complexes were supplemented with 1.7 mM deoxyribonucleoside triphosphates (dATP, dGTP, dCTP, rTTP), 1.0 μL labelled primer, 33 mM MgCl_2_, 5U AMV-RT (Promega), 80U RiboLock (Thermo Fisher) and incubated (37°C, 45 min) prior to addition of addition of 100 μL 25 mM EDTA and 0.5 w/v % SDS to stop the reaction. Samples were phenol:chloroform extracted, ethanol precipitated and resuspended in 10 μL loading dye (0.05 w/v % bromophenol blue, 0.05 w/v % xylene cyanol, 20 mM EDTA pH 8.0, 91 % formamide).

Sanger sequencing ladders were generated as previously described (Pisarev *et al*., 2007). Briefly, 1.0 μg primer was combined with 3.0 μg plasmid DNA template in 20 μL water, prior to addition of NaOH (0.2 M). Annealing mixtures were incubated (5 min), diluted with water, ethanol precipitated and resuspended in sequencing buffer (40 mM Tris-HCl pH 7.5, 20 mM MgCl_2_, 50 mM NaCl). Separately, four ddNTP mixes were prepared containing 80 μM dNTPs and 8.0 μM of either ddATP, ddTTP, ddCTP or ddGTP. Sequencing reactions were assembled as directed by the manufacturer (Sequenase, ThermoFisher) and incubated at 45 °C for 10 min prior to addition of loading dye.

Primer extension samples were run alongside sequencing reactions on a 6.0 % Urea-PAGE gel (40 W constant, 2 h). Gels were dried under vacuum (80°C, 2 h) and analysed by autoradiography using a phosphor storage plate and Typhoon 5 scanner (Amersham).

#### Band quantification

Efficiency of 48S complex formation was estimated through band quantification of autoradiographs in duplicate using Image Studio Lite (v6.1.0.79). Local background was estimated using the median method within a 3-pixel border surrounding each band. This was subtracted from the signal corresponding to the 48S toeprint bands and the control bands. Where indicated by an ‘*’ on gel images, a characteristic band corresponding to the edge of domain V was used as a control to account for differences in reverse transcriptase activity, cDNA recovery or sample loading. The total signal for each 48S toeprint band was divided by that of the corresponding control band in the same lane. These ratios were then expressed relative to a positive control sample run on the same gel (e.g. the unmutated IRES), which was arbitrarily defined as having 100% activity. Quantification was carried out independently from two different gel images, each originating from an independent reconstitution experiment.

#### Cryo-EM sample preparation

Quantifoil grids (R2/2, 300 mesh, copper) were glow discharged using a Harrick Plasma cleaner set on High (30W) for 60s at 0.6 Torr. 48S complexes were assembled on the PV-AUG_586_-good IRES by mixing 0.26 μM 40S with 0.5 μM eIFs, 0.5 μM PCBP2, 1 μM IRES RNA and 29 μM total tRNA (pre-treated with aa-tRS to charge initiator tRNA) in buffer (20 mM Tris-HCl pH 7.5, 60 mM KCl, 2.5 mM MgCl_2_, 2.0 mM DTT, 0.25 mM spermidine, 0.5 mM GTP, 2.0 mM ATP). Samples were incubated (37°C, 15 min) before application of 2.0 μL onto grids using a Vitrobot Mark IV (Thermo Scientific). Grids were blotted for 2-4 s (100% humidity, -10 force) before plunging into liquid ethane.

#### Cryo-EM data acquisition and image processing

Grids were screened on a 200 kV Glacios electron cryo-microscope equipped with a Falcon 4 detector (Thermo) at the York Structural Biology Laboratory, University of York, UK. Subsequently, datasets were acquired on a 300 kV Titan Krios electron cryo-microscope equipped with a K3 detector and Gatan Imaging Filter located at the electron Bio-Imaging Centre (eBIC), Diamond Light Source. A total of 26,797 micrograph movies were recorded in aberration-free image shift mode using defocus targets of -2.2, -2.0, -1.8, -1.6, -1.4, -1.2, and -1.0 μm. Per movie, a total dose of 50 e^-^/Å^2^ was applied over 2.07 s. A nominal magnification of 105,000 was applied resulting in a final uncalibrated object sampling of 0.831 Å pixel size. Data collection parameters are detailed in **Appendix Table S1**.

The cryo-EM datasets were processed using RELION 5.0 (Scheres, 2012) as summarised in **Figures EV3 and EV4**. Motion correction with dose-weighting was performed with 5 x 5 patches using the RELION implementation of UCSF MotionCor2 (Zheng *et al*, 2017), prior to CTF estimation with CTFFIND 4.1 (Rohou & Grigorieff, 2015). A Topaz model (Bepler *et al*, 2019) was used to pick 3,029,414 particles. Particle images were initially downscaled four-fold and extracted in a 120-pixel box (3.32 Å effective pixel size). Particles were subjected to three rounds of two-dimensional classification, resulting in 612,557 high-quality 40S-containing particles. These particles were used to generate a consensus 6.7 Å map using an initial 3D reference of a canonical 48S initiation complex low-pass filtered to 80 Å (Brito Querido *et al*., 2020). Several rounds of 3D focused classification were then performed, implemented as masked 3D classification without alignment jobs in RELION 5.0. In each case, the T-value was optimised *de novo* based on the masked volume, and after a successful separation, angular assignments were updated with masked 3D refinement. In short, particles were separated first on occupancy of eIF3, then tRNA, eIF2 and 40S head rotation, and finally the presence and positioning of IRES dIVc.

Particles were first separated through focused classification of eIF3 which yielded three subsets. Subset 1 (296,351 particles) contained ribosomes lacking eIF3, but with eIF2, tRNA, eIF1A, and IRES dIVc present. Subset 2 (88,275 particles) contained ribosomes with eIF3, eIF2, tRNA, eIF1A, and IRES dIVc. Subset 3 (226,646 particles) only contained 40S and eIF1A and was not processed further. Subset 2 underwent further focused classification based on eIF2 and 40S head rotation state. This yielded **structure 1**, an ‘open’ scanning 48S complex (10,518 particles), a ‘closed’ 48S complex at AUG_586_ (23,333 particles) and 40S-eIF3-eIF1A, which was not processed further.

Next, these closed 48S complexes from subset 2 were subjected to further focused classification based on the presence of IRES dIVc (11,241 particles). Subset 1 underwent multiple rounds of focused classification based on dIVc, and the resultant particles were combined with the dIVc-containing particles from subset 2. This set of particles (44,492) has maximum dIVc occupancy and was used for **structure 2** but is sub-stoichiometric with respect to eIF3. Particles from structure 1 and structure 2 were re-extracted to the nominal pixel size of 0.831 with subsequent 3D refinement yielding resolutions of 4.0 Å and 3.2 Å, respectively. Further improvement was achieved by applying CTF refinement for beam tilt, trefoil, fourth order aberration and magnification anisotropy. CTF was estimated per particle, and astigmatism was estimated per micrograph. 3D refinement of **structures 1 and 2**, with subsequent gold-standard FSC estimation yielded resolutions of 3.8 Å and 2.9 Å, respectively (**Appendix Figures S2 and S3**). Upon inspection of the structure 2 maps, conformational heterogeneity was apparent for the dIVc_1_ segment. We therefore decided to perform masked classification without angular searches, focused on dIVc. This produced **structures 3 and 4** with altered positions of dIVc_1_. After 3D refinement and gold-standard FSC estimation, the reported resolutions for structures 3 and 4 are 3.1 Å and 3.6 Å, respectively (**Appendix Figures S4 and S5**). Finally, maps weighted by estimated local resolution were calculated.

#### Model building and refinement

As a starting point, atomic models of human 48S initiation complexes and sub-components were downloaded from the wwPDB (‘open’ scanning 48S, Data ref: PDB 6ZMW, 2020; ‘closed’ 40S, Data ref: PDB 7TQL, 2022; eIF2 and tRNA, Data ref: PDB 6YBV, 2020; tRNA and eIF1A, Data ref: PDB 8PJ2, 2023. Chains corresponding to mRNA were removed, and all chains renumbered. ChimeraX (Meng *et al*, 2023) was first used for manual positioning and rigid-body fitting of the models into unsharpened cryo-EM maps, prior to per-chain rigid-body adjustments using ‘jiggle fit’ in COOT (Casañal *et al*, 2020). For flexible fitting of individual subunits into weaker density, real space ‘chain refine’ was used with maps weighted by local resolution. For this step, a network of all-chain self-restraints was generated to preserve backbone and side-chain geometry (5.0 Å distance, German-McClure alpha 0.01-0.3, depending on local map quality).

An initial model of PV IRES dIVc was generated by AlphaFold3 (Abramson *et al*, 2024), comprising three segments corresponding to dIVc_1_, dIVc_2_ and dIVc_3_. Initial rigid-body docking into the map was guided by the tRNA-GNRA contact (dIVc_3_) for which the most characteristic density was observed, including the positioning of the single-stranded RNA loop. However, the angles between helical segments required remodelling. To facilitate this at low resolution whilst preserving RNA geometry, chain breaks were manually introduced in the linker regions between dIVc_1_/IVc_2_ and dIVc_2_/IVc_3_, creating single-stranded RNA ‘hinges’ about which restrained helical segments could move independently. All-atom self-restraints were generated (4.3 Å and 3.7 Å distances) to preserve base pairing, stacking and base-pair planarity during subsequent ‘chain refine’ operations, guided by a local resolution-weighted map. Finally, the chain breaks were manually rebuilt.

In all cases, atomic model refinement was carried out using Servalcat (Yamashita *et al*, 2021) in CCP-EM (Burnley *et al*, 2017) via a Doppio wrapper. Unfiltered half-maps were used as the refinement target, with a maximum resolution cutoff corresponding to the FSC at 0.143. To stabilise refinement of chains in lower-resolution areas, jelly-body restraints (4.2 Å distance, sigma 0.02) were applied, and a manual experimental data vs. stereochemistry weighting of 0.5 was selected. Hydrogens were refined in the ‘riding’ position. MolProbity (Williams *et al*, 2018) was used to assess geometry after each cycle, and models were manually corrected in COOT, minimising clashes, bad rotamers and Ramachandran outliers. Structural figures were rendered in PyMol (Schrödinger, LLC) and ChimeraX (Meng *et al*., 2023). Model refinement statistics are detailed in **Appendix Table S1**.

#### SHAPE analysis

SHAPE was performed as previously described (Wilkinson *et al*, 2006). 24 pmol of *in vitro* transcribed and purified RNA (full-length WT PV 5′ UTR) was heat denatured at 95°C for 2 min followed by snap cooling on ice for 2 min. RNA was refolded at 37°C for 30 min in folding buffer (100 mM HEPES, 6.0 mM MgCl_2_ and 100 mM KCl final concentration). For RNA+PCBP2 experiments, refolded RNA was further incubated with a molar excess of PCBP2 (3.0 µM) for 10 min at 37°C. PCBP2 was incubated at 30°C for 10 min with Ribolock RNase inhibitor (40 U/µl, Thermo) *<u>before</u>* adding to the refolded RNA. Refolded RNA was divided in equal volumes into separate tubes and treated with either NMIA (*N*-methylisatoic anhydride) dissolved in DMSO (10 mM final concentration) or DMSO only for 50 min at 37°C. RNA was mixed with 100 μL of EDTA (100 mM), 4.0 μL of NaCl (5.0 M) and 2.0 μL of glycogen (20 mg/mL), ethanol precipitated and resuspended in 30µl 0.5xTE buffer. For primer extension, RNA was mixed with 5′ VIC labelled primer (1.5-2.0 µM final concentration, Applied Biosystems). Labelled primer was annealed to the modified RNA at 65°C for 10 min and 35°C for 5 min followed by reverse transcription as per the manufacturer’s instructions using

Superscript III (Invitrogen) at 52°C for 30 min. For sequencing ladders, 24 pmol of untreated RNA was reverse transcribed in the presence of ddCTP (10 mM, Jena Biosciences), using the above primer labelled with 5′ NED. All primer extensions were terminated by alkaline hydrolysis, neutralised with HCl and ethanol precipitated before being resuspended in ddH_2_O. cDNAs from NMIA- and DMSO-treated RNA were mixed with equal amounts of sequencing ladder and final samples were dried onto sequencing plates before resuspension in formamide followed by fractionation through capillary electrophoresis (Applied Biosystems 3130xl analyser, University of Cambridge, Department of Biochemistry DNA Sequencing Facility). SHAPE reactivities were calculated in QuSHAPE software (Karabiber *et al*, 2013) using default parameters.

#### Cells

HeLa cells (ATCC) were maintained at 37°C in Dulbecco’s Modified Eagle Medium (DMEM, Lonza) supplemented with 10% fetal bovine serum (FBS), 1.0 mM L-glutamine, 20 mM HEPES (pH 7.3), and Penicillin/Streptomycin (10,000 U/mL). HIEC6 cells (human intestinal epithelial cell line, isolated from a healthy donor, ATCC, CRL-3266) were maintained in Opti-MEM (Gibco) containing 20 mM HEPES, 1.0 mM L-Glutamine, Penicillin/Streptomycin (10,000 U/mL), 10 ng/mL hEGF (Sigma-Aldrich), and 5% FBS. All cells were tested mycoplasma negative throughout the work (MycoAlert^®^ Mycoplasma Detection Kit, Lonza).

#### Plasmids

To evaluate the efficiency of IRES-dependent translation, a previously described dual-luciferase reporter plasmid containing a T7 promoter, cap-dependent Renilla luciferase gene followed by the CVA13 5′ UTR sequence, a 2A StopGo sequence, then the firefly luciferase gene and CVA13 3′ UTR sequence (O’Connor *et al*., 2026). All mutations were introduced using site-directed mutagenesis and confirmed by Sanger sequencing. The resulting plasmids were linearized with NdeI before T7 RNA transcription. The CVA13 replicon was designed using previously described reverse genetics clone (O’Connor *et al*., 2026). Briefly, the capsid-encoding sequence was replaced by the mCherry gene while preserving the original polyprotein cleavage sites. The resulting clone was sequenced using the Plasmidsaurus service and deposited in GenBank (accession number PX114590). All mutations were introduced using site-directed mutagenesis and confirmed by Sanger sequencing. The resulting plasmids were linearized with EagI before T7 RNA transcription.

#### RNA transcription for reporter and replicon assays

Linearized replicon-encoding plasmids were used as templates to generate RNAs using Jena HighYield T7 ARCA mRNA Synthesis kit according to manufacturer’s instructions. The replicon RNA was purified using Zymo RNA Clean & Concentrator kit and quantified.

#### IRES reporter assays

HeLa and HIEC6 cells were transfected in triplicate using the method previously described (Lulla *et al*., 2019). For each transfection, 100 ng of purified T7 RNA combined with 1.0 µl Lipofectamine 2000 (Invitrogen) in 10 µl Opti-Mem (Gibco) supplemented with RNaseOUT (Invitrogen; diluted 1:1,000 in Opti-Mem) were added to each well containing 6×10^4^ cells on a 96-well plate. Transfected cells were supplemented with 5% FBS and incubated at 37°C for 8 hours. Cells were lysed in 100 µl Passive Lysis Buffer (Promega) and freeze-thawed. Luciferase activity was measured using the Dual Luciferase Stop & Glo Reporter Assay System (Promega) as per manufacturer’s instructions. IRES activity was calculated as the ratio of IRES-dependent translation (firefly) to cap-dependent translation (Renilla). Four independent experiments were performed to confirm the reproducibility of the results.

#### Enterovirus replicon assays

HeLa and HIEC6 cells (6×10^4^ cells on a 96-well plate) were transfected in quadruplicate using the above-described reverse transfection protocol and previously validated controls (Ali *et al*, 2024). Guanidine hydrochloride (GuHCl) was included as a replication-deficient control at a concentration of 2.0 mM. The mCherry signal was assessed using live cell imaging with the Incucyte SX5, an automated phase-contrast and fluorescence microscope, within a humidified incubator. At 1.5-hour intervals, a single whole-well image was taken and used to measure mCherry integrated intensity using SX5-integrated software. The mCherry signal level peaked at 12 hours post-transfection, and this time point was used to evaluate replication efficiency. Four independent experiments were performed to confirm the reproducibility of the results.

## Expanded View Figures and Legends

**Figure EV1.**
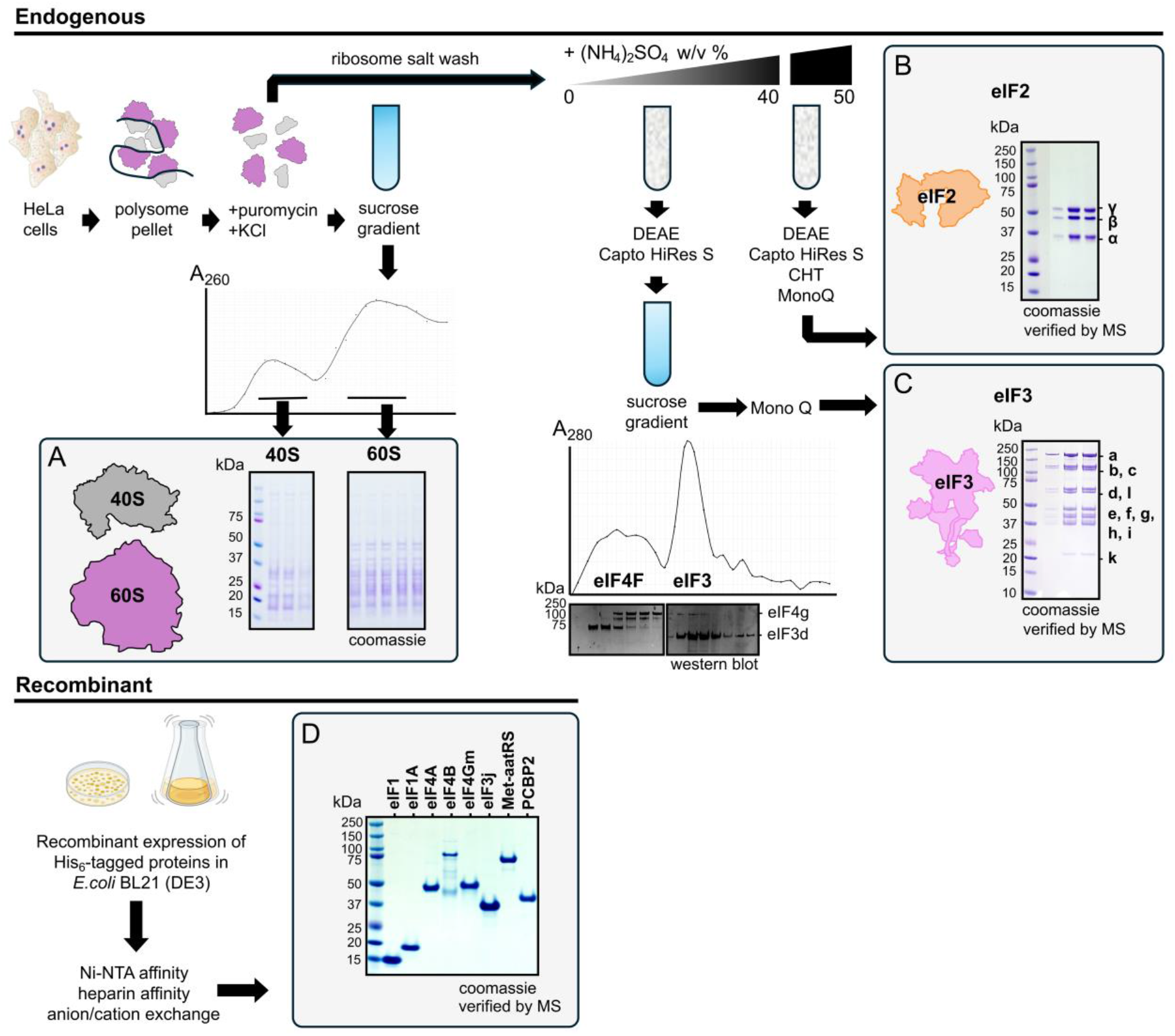
**Scheme for the purification of human ribosome subunits and initiation factors**, starting from either HeLa cells *(upper)* or recombinant protein expression *(lower)*. Proteins were analysed by SDS-PAGE and visualised by Coomassie staining. All purified proteins were verified by mass spectrometry. **A,** 40S and 60S ribosomal subunits are purified via sucrose gradients following puromycin and salt treatment of polysomes isolated from HeLa cytoplasm. The ribosome salt wash fraction is then dialysed and fractionated by progressive ammonium sulfate precipitation. **B,** The 40-50% fraction is used to purify endogenous eIF2. **C,** The 0-40 % fraction is used to purify endogenous eIF3 and eIF4F. **D,** eIFs 1, 1A, 4A, 4B, 4Gm, 3j, the Methionyl aminoacyl tRNA synthetase (Met-aatRS) and poly-C binding protein (PCBP2) are purified following recombinant expression in *E.coli* BL21.

**Figure EV2.**
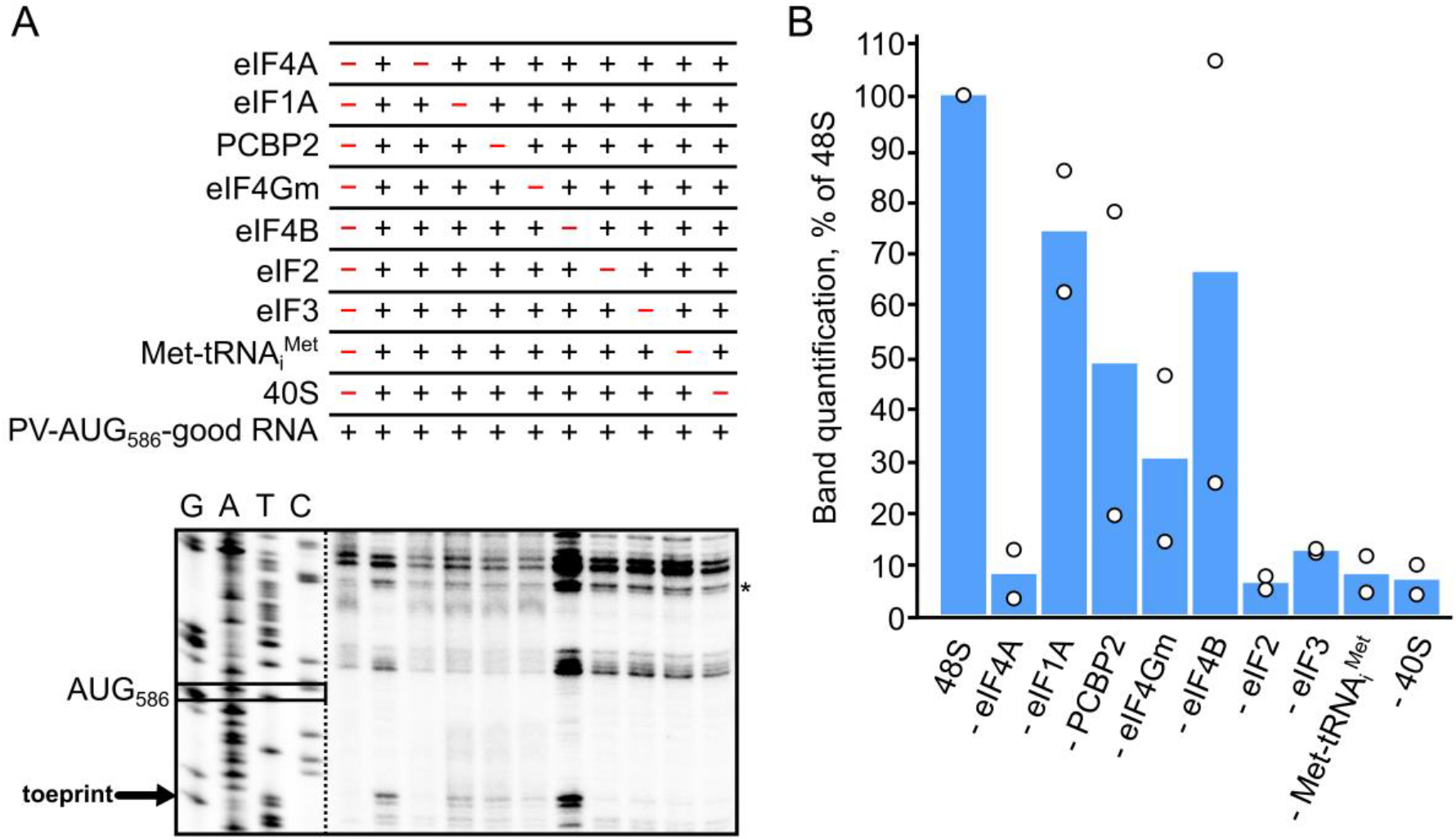
Effect of factor omission on 48S complex formation on AUG_586_ on the PV-AUG_586_-good IRES A, Analysis of primer extension assay by Urea-PAGE and autoradiography, demonstrating the effects of omission of factors on human 48S complex formation at AUG_586_ on the PV-AUG_586_-good IRES. ‘*’ Denotes the band used for normalization. The expected position of toeprints corresponding to AUG_586_ is indicated (black arrow). Sanger sequencing ladder is displayed at a different contrast for ease of reading sequence. **B,** Band quantification showing the effect of human 48S complex formation at AUG_586_ on the PV-AUG_586_-good IRES upon omission of different factors, relative to a control which includes all required factors. (n=2 technical replicates, bar represents mean, individual data points are shown, replicate data in **Appendix Figure S1A**). See **Methods and Protocols** for details of band quantification.

**Figure EV3.**
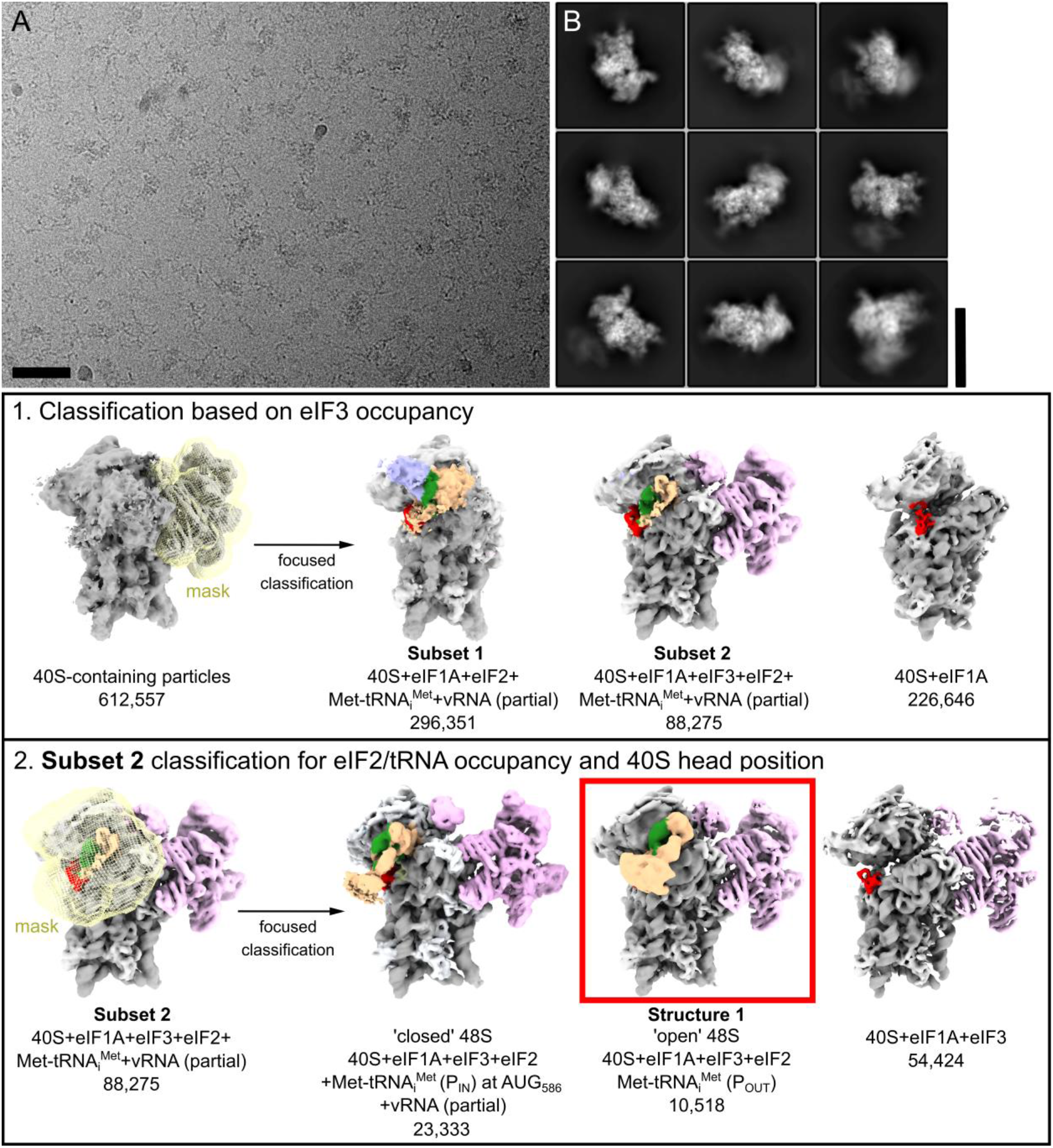
Summary of cryo-EM processing and particle classification. **A,** Representative electron cryo-micrograph, acquired on a Titan Krios microscope with a K3 camera (105,000 x magnification; scale bar = 50 nm). **B,** Representative 2D class averages of 40S-containing particles (scale bar = 23 nm). Compositional and conformational heterogeneity is apparent. Details of the key steps in particle classification. Intermediate cryo-EM maps (3.32 Å / px; 3.0 **–** 5.0 σ) are colour-coded according to component identity as in Figure 3A. Masks used for focused classification are displayed as meshes superimposed on the parent map. The composition and number of particles in each class is indicated.

**Figure EV4.**
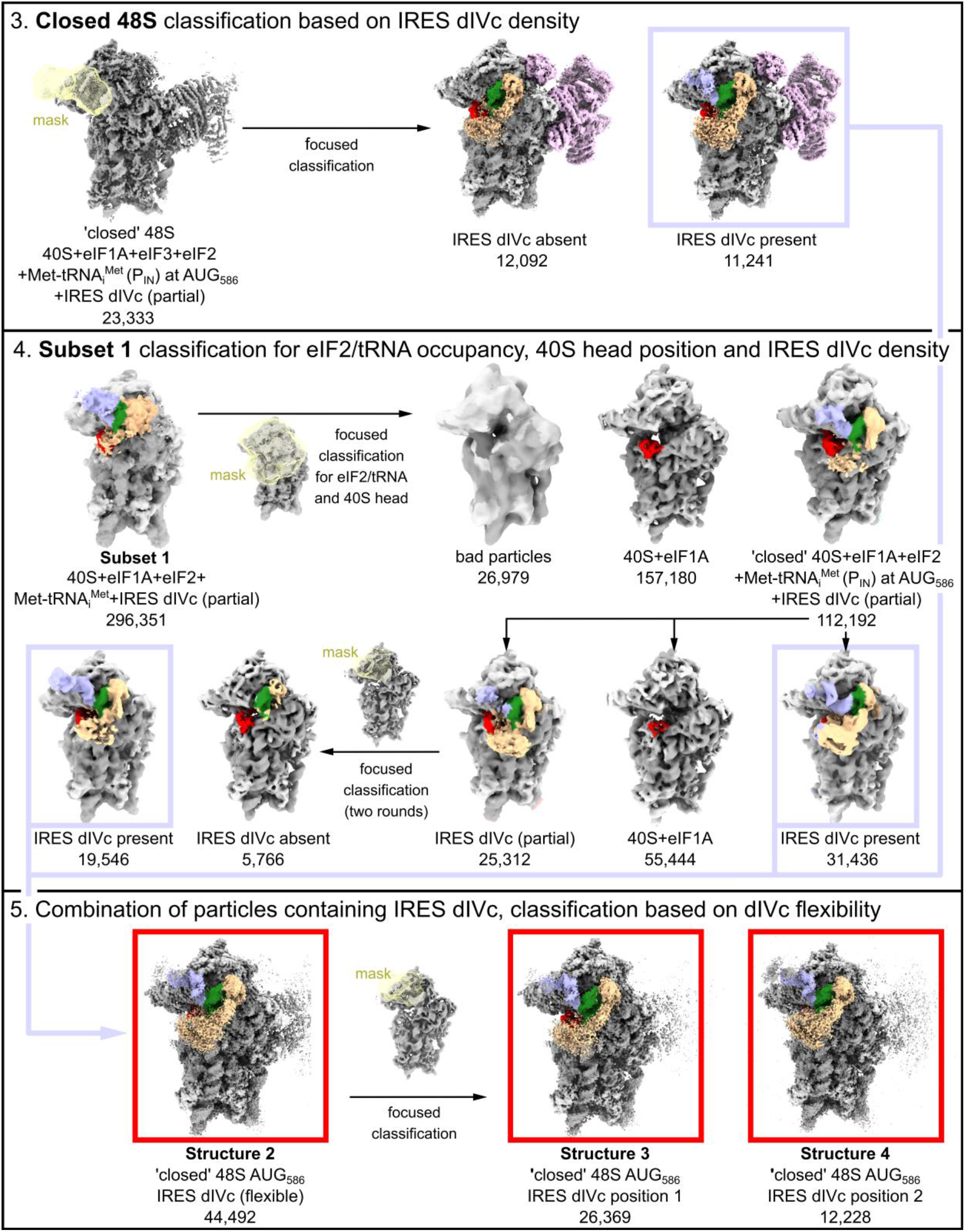
Summary of cryo-EM processing and particle classification. Details of the key steps in particle classification. Intermediate cryo-EM maps, masks and class annotations are displayed as above. Final maps (0.831 Å / px; 3.0 σ) are boxed in red.

**Figure EV5.**
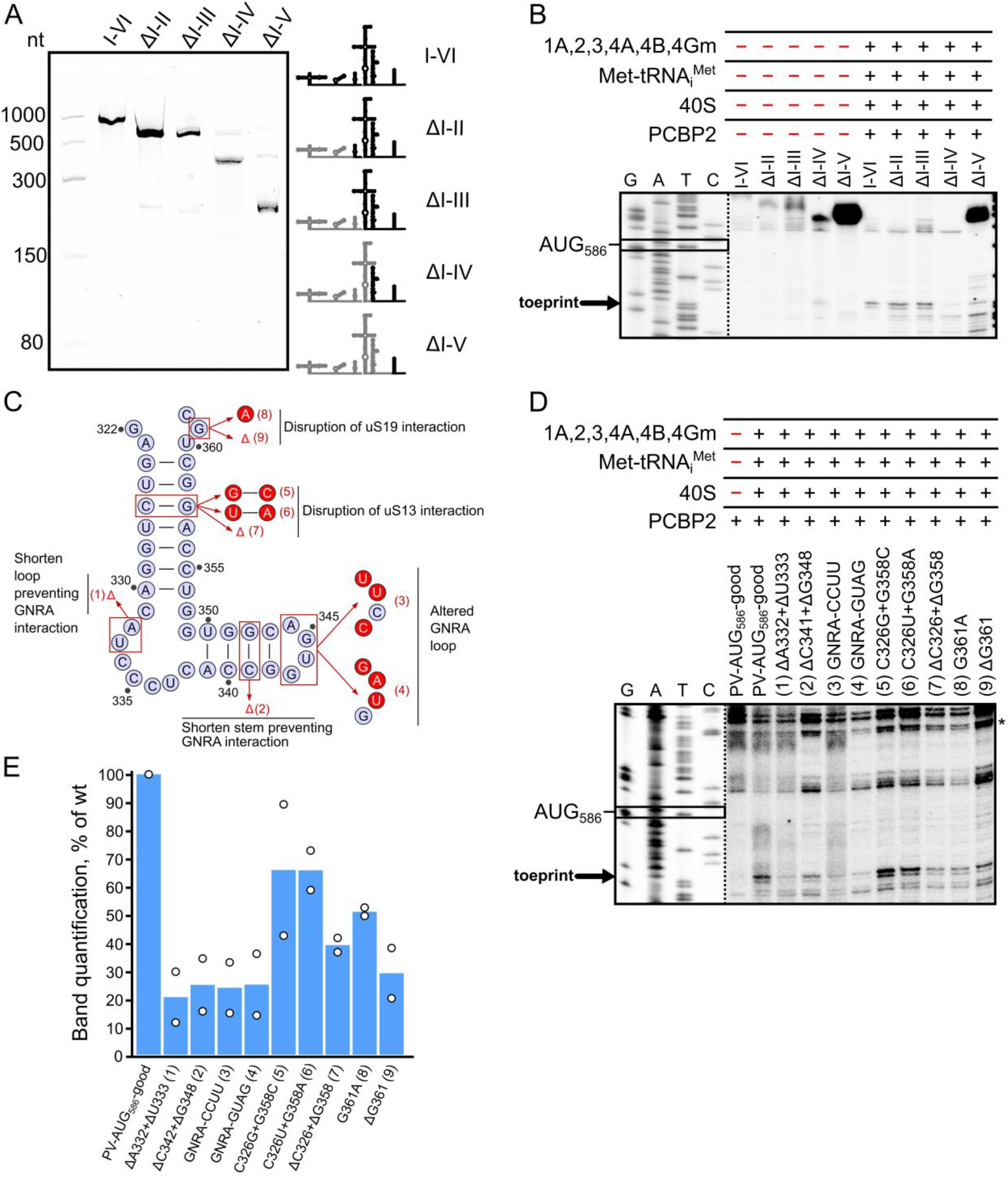
dIV is required for efficient 48S complex formation on PV-AUG_586_-good IRES A, *(Left)* Urea-PAGE analysis of domain truncations of PV-AUG_586_-good IRES, following *in vitro* transcription. *(Right)* Schematic diagram illustrating effects of truncations on the IRES. **B,** Analysis of primer extension assay by Urea-PAGE and autoradiography, demonstrating the effects of domain truncation on human 48S complex formation at AUG_586_ on the PV-AUG_586_-good IRES. Sanger sequencing ladder is displayed at a different contrast for ease of reading sequence (n=2, see technical replicate in **Appendix Figure S1B**). **C,** Location of mutations on the PV dIVc sequence are indicated (red). **D,** Analysis of primer extension assay by Urea-PAGE and autoradiography, demonstrating the effects of dIVc mutations on human 48S complex formation at AUG_586_ on the PV-AUG_586_-good IRES. ‘*’ Denotes the band used for normalization. Sanger sequencing ladder is displayed at a different contrast for ease of reading of sequence. **E,** Band quantification showing the effect of human 48S complex formation at AUG_586_ on the PV-AUG_586_-good IRES upon mutation of dIVc. (n=2 technical replicates, bar represents mean, individual data points are shown, replicate data in **Appendix Figure S1C**). See **Methods and Protocols** for details of band quantification.

## Data Availability

Cryo-EM maps, half-maps and masks for structures 1-4 have been deposited to the EMDB under accession numbers EMD-59031 (structure 1), EMD-59032 (structure 2), EMD-59033 (structure 3) and EMD-59034 (structure 4). Refined atomic models for structures 1-4 have been deposited to the wwPDB under accession numbers 32OB/pdb_000032ob (structure 1), 32OC/pdb_000032oc (structure 2), 32OE/pdb_000032oe (structure 3) and 32OG/pdb_000032og (structure 4). **Prior to public release, deposited coordinates, maps and validation reports are available on Google Drive**: (https://drive.google.com/drive/folders/13UzDBktpbfmyaCB-W57-LbdwPyUSHe2d?usp=sharing)

The CVA13 replicon sequence has been deposited in GenBank (accession number PX114590). Several deposited cryo-EM structures were used for analysis or comparisons, as follows. Data ref: PDB 8PJ1, 2024; Data ref: PDB 7QP6, 2022; Data ref: PDB 6ZMW, 2020; Data ref: PDB 8G5Y, 2023; Data ref: PDB 8PJ2, 2024; Data ref: PDB 8PJ3, 2024; Data ref: PDB 8PJ4, 2024; Data ref: PDB 8PJ5, 2024; Data ref: PDB 7TQL, 2022; Data ref: PDB 6YBV, 2020; Data ref: PDB 8SUP, 2026; Data ref: PDB 9UZK, 2026; Data ref: PDB 9UZL, 2026

## Acknowledgements

We thank Sam Hart and Johan Turkenburg in the YSBL cryo-EM facility for technical assistance, and we acknowledge the support of the Wolfson Foundation, Wellcome and BBSRC (BB/Z515668/1). We thank Michael Plevin and Michelle Hawkins for useful scientific discussions. We acknowledge Diamond Light Source for access to a Titan Krios microscope at the UK national Electron Bio-imaging Centre (eBIC) under proposal BI34172, funded by the Wellcome Trust, MRC and BBRSC. The Viking cluster was used during this project, which is a high-performance compute facility provided by the University of York. We are grateful for computational support from the University of York, IT Services and the Research IT team. M.A.A.V is supported by a BBSRC PhD studentship from the White Rose Mechanistic Biology DTP (BB/M011151/1). J.H., K.F., and V.L. are supported by an MRC project grant (MR/T000376/1) and a Sir Henry Dale Fellowship (220620/Z/20/Z) from the Wellcome Trust and the Royal Society to V.L. S.S.N. and T.R.S. are supported by a Sir Henry Dale Fellowship (202471/Z/16/A) from the Wellcome Trust and the Royal Society to TS, and BBSRC grants BBS/E/PI/230002A and BBS/E/PI/23NB0003 to The Pirbright Institute. C.H.H. is supported by a Sir Henry Dale Fellowship (221818/Z/20/Z) from the Wellcome Trust and the Royal Society, and by the Lister Institute of Preventive Medicine. For the purpose of open access, the authors have applied a CC-BY public copyright licence to any Author Accepted Manuscript version arising from this submission.

