## Appendix (Supplementary Material) for "Structural and mechanistic insights into translation initiation on the enterovirus Type 1 IRES"

### Structural Data

Cryo-EM maps, half-maps, masks and refined atomic coordinates have been deposited to the wwPDB and EMDB, under the following accession numbers:

**Structure 1 'open' 48S**........ **PDB** 32OB/pdb_000032ob; **EMDB** EMD-59031

**Structure 2 'closed' 48S at $AUG_{586}$**........ **PDB** 32OC/pdb_000032oc; **EMDB** EMD-59032

**Structure 3 'closed' 48S at $AUG_{586}$, $dIVc_1$ position 1**........ **PDB** 32OE/pdb_000032oe; **EMDB** EMD-59033

**Structure 4 'closed' 48S at $AUG_{586}$, $dIVc_1$ position 2**........ **PDB** 32OG/pdb_000032og; **EMDB** EMD-59034

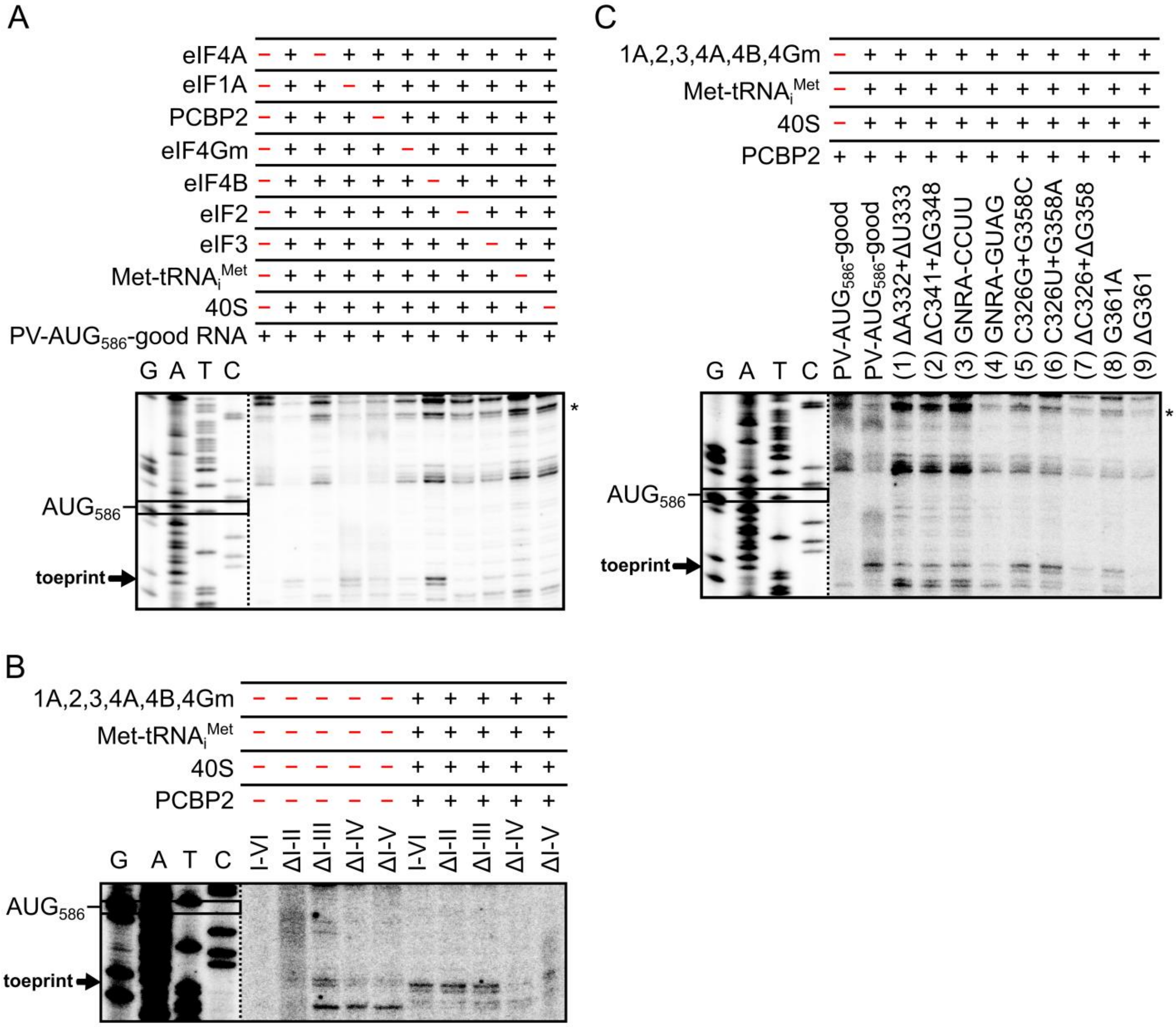

**Appendix Figure S1 – Replicate autoradiographs of 48S complex formation for dIVc mutants, factor omission and domain truncation experiments**

**A,** Analysis of primer extension assay by Urea-PAGE and autoradiography, demonstrating the effects of omission of factors on human 48S complex formation at $AUG_{586}$ on the PV-$AUG_{586}$-good IRES. '*' Denotes the band used for normalization. Sanger sequencing ladder is displayed at a different contrast for ease of reading of sequence. Replicate experiment for **Figure EV2A.**

**B,** Analysis of primer extension assay by Urea-PAGE and autoradiography, demonstrating the effects of domain truncation on human 48S complex formation at $AUG_{586}$ on the PV-$AUG_{586}$-good IRES. Replicate experiment for **Figure EV5B.**

**C,** Analysis of primer extension assay by Urea-PAGE and autoradiography, demonstrating the effects of dIVc mutations on human 48S complex formation at $AUG_{586}$ on the PV-$AUG_{586}$-good IRES. '*' Denotes the band used for normalization. Sanger sequencing ladder is displayed at a different contrast for ease of reading of sequence. Replicate experiment for **Figure EV5D.**

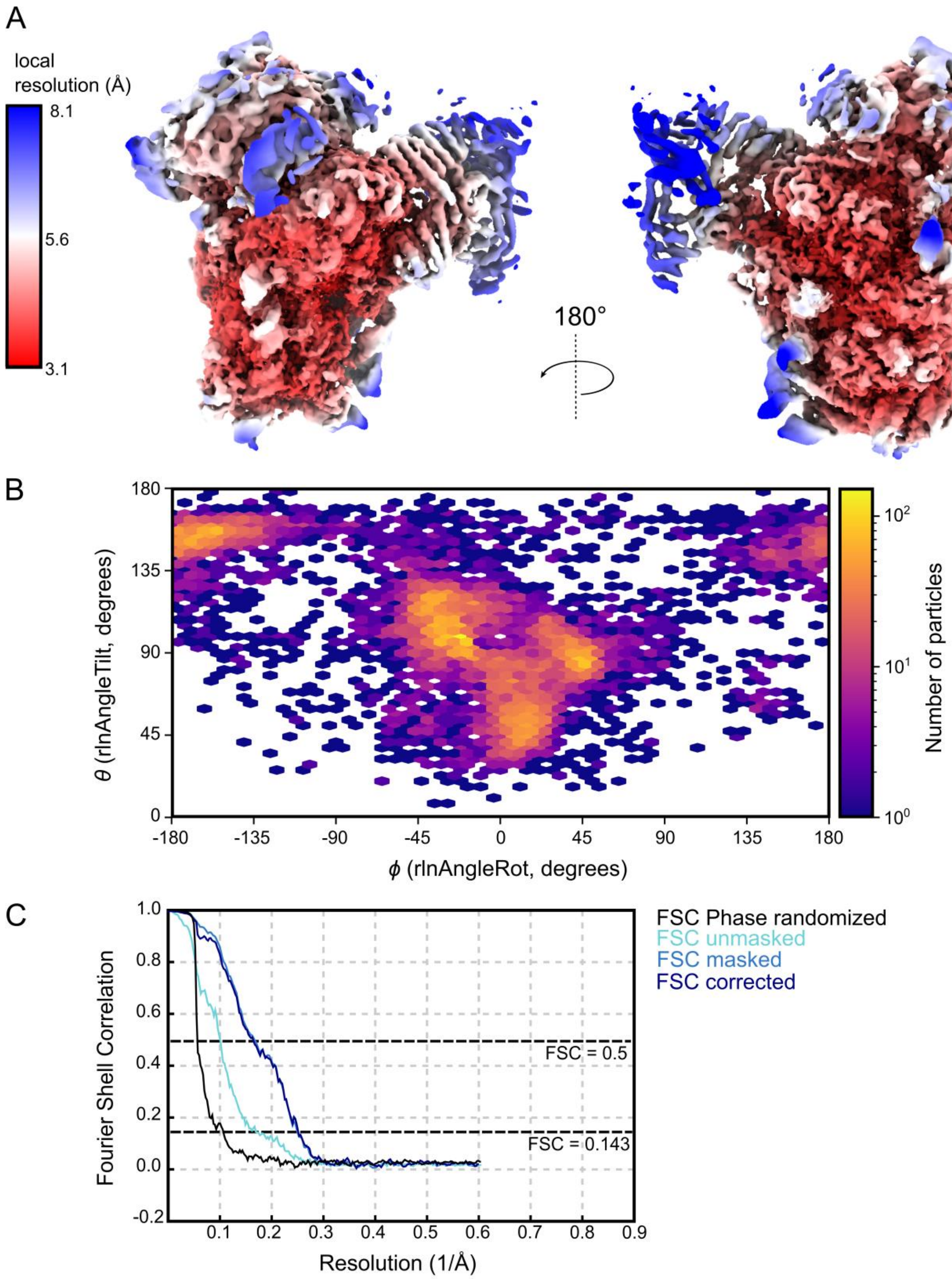

**Appendix Figure S2 - Validation of the cryo-EM map for structure 1 ('open' 48S complex)**

**A,** Local-resolution weighted cryo-EM map for 'open' 48S complex (0.831 Å / px, contoured at 3.3 σ), coloured by estimated local resolution as indicated by the heatmap key.

**B,** Euler angle distribution plot of particles in the final reconstruction.

**C,** Gold-standard Fourier shell correlation (FSC) curves for final map generated by RELION post-processing. Masked (light blue), unmasked (cyan), and phase randomized (black) are shown.

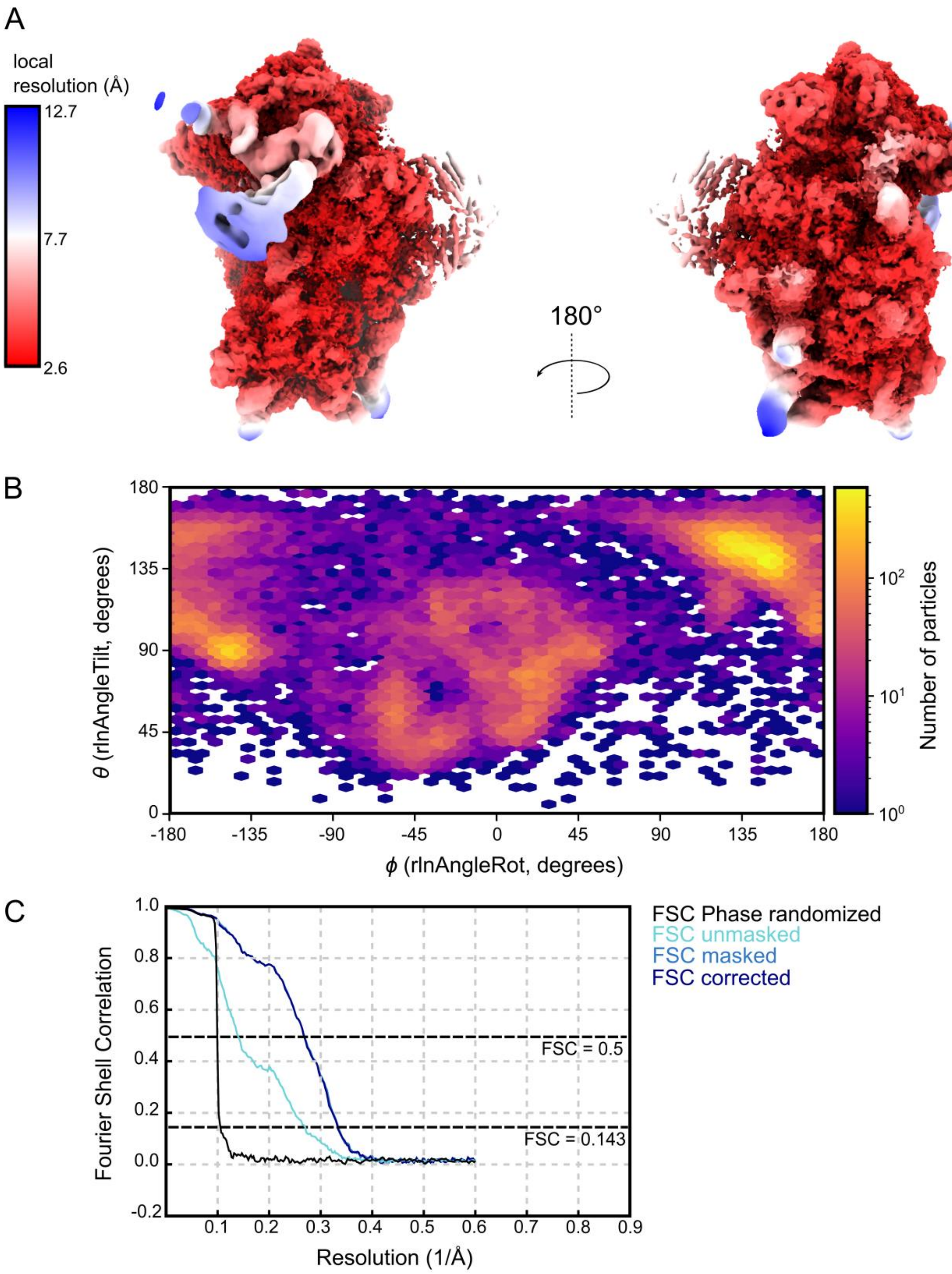

**Appendix Figure S3 - Validation of the cryo-EM map for structure 2 ('closed' 48S complex)**

**A,** Local-resolution weighted cryo-EM map for 'closed' 48S complexes at $AUG_{586}$ (0.831 Å / px, contoured at 2.0 σ), coloured by estimated local resolution as indicated by the heatmap key.

**B,** Euler angle distribution plot of particles in the final reconstruction.

**C,** Gold-standard Fourier shell correlation (FSC) curves for final map generated by RELION post-processing. Masked (light blue), unmasked (cyan), and phase randomized (black) are shown.

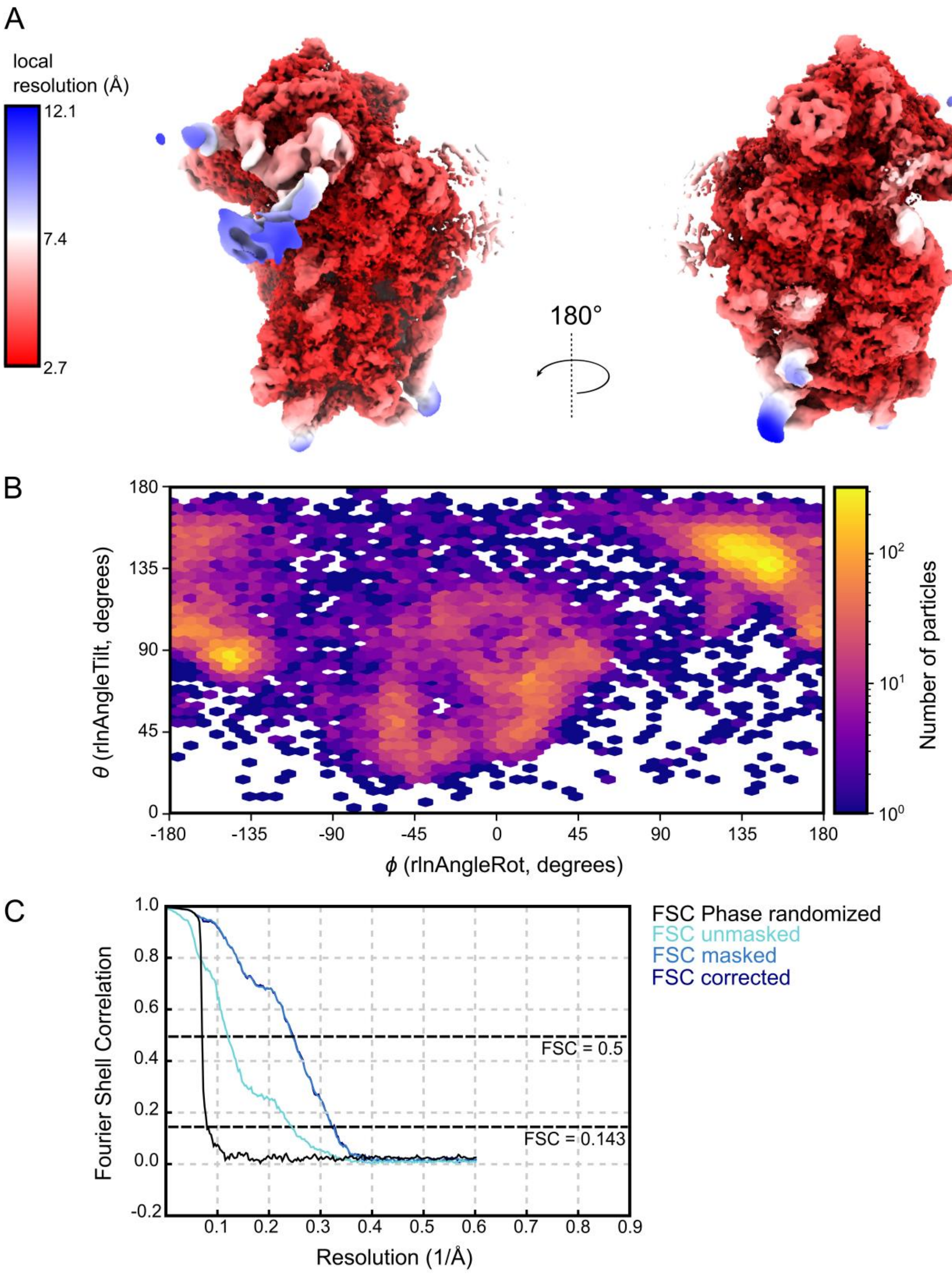

**Appendix Figure S4 - Validation of the cryo-EM map for structure 3 (dIVc$_1$ position 1)**

**A,** Local-resolution weighted cryo-EM map for 'closed' 48S complexes at AUG$_{586}$ with dIVc$_1$ in position 1 AUG$_{586}$ (0.831 Å / px, contoured at 2.2 σ), coloured by estimated local resolution as indicated by the heatmap key.

**B,** Euler angle distribution plot of particles in the final reconstruction.

**C,** Gold-standard Fourier shell correlation (FSC) curve for final map generated by RELION post-processing. Masked (light blue), unmasked (cyan), and phase randomized (black)

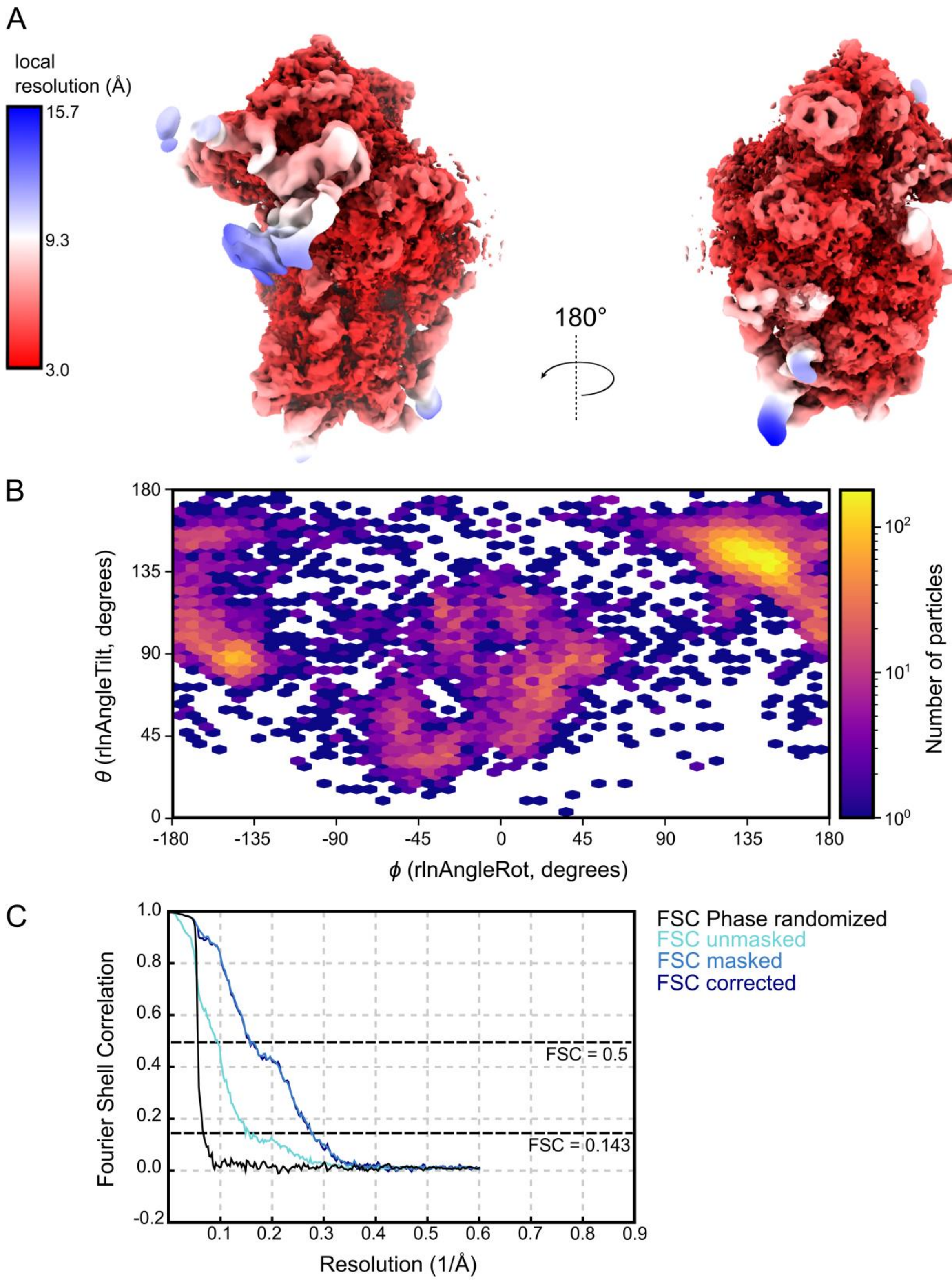

**Appendix Figure S5 - Validation of the cryo-EM map for structure 4 ($dIVc_1$ position 2)**

**A,** Local-resolution weighted cryo-EM map for ‘closed’ 48S complexes at $AUG_{586}$ with $dIVc_1$ in position 2 $AUG_{586}$ (0.831 Å / px, contoured at 2.5 σ), coloured by estimated local resolution as indicated by the heatmap key.

**B,** Euler angle distribution plot of particles in the final reconstruction.

**C,** Gold-standard Fourier shell correlation (FSC) curve for final map generated by RELION post-processing. Masked (light blue), unmasked (cyan), and phase randomized (black)

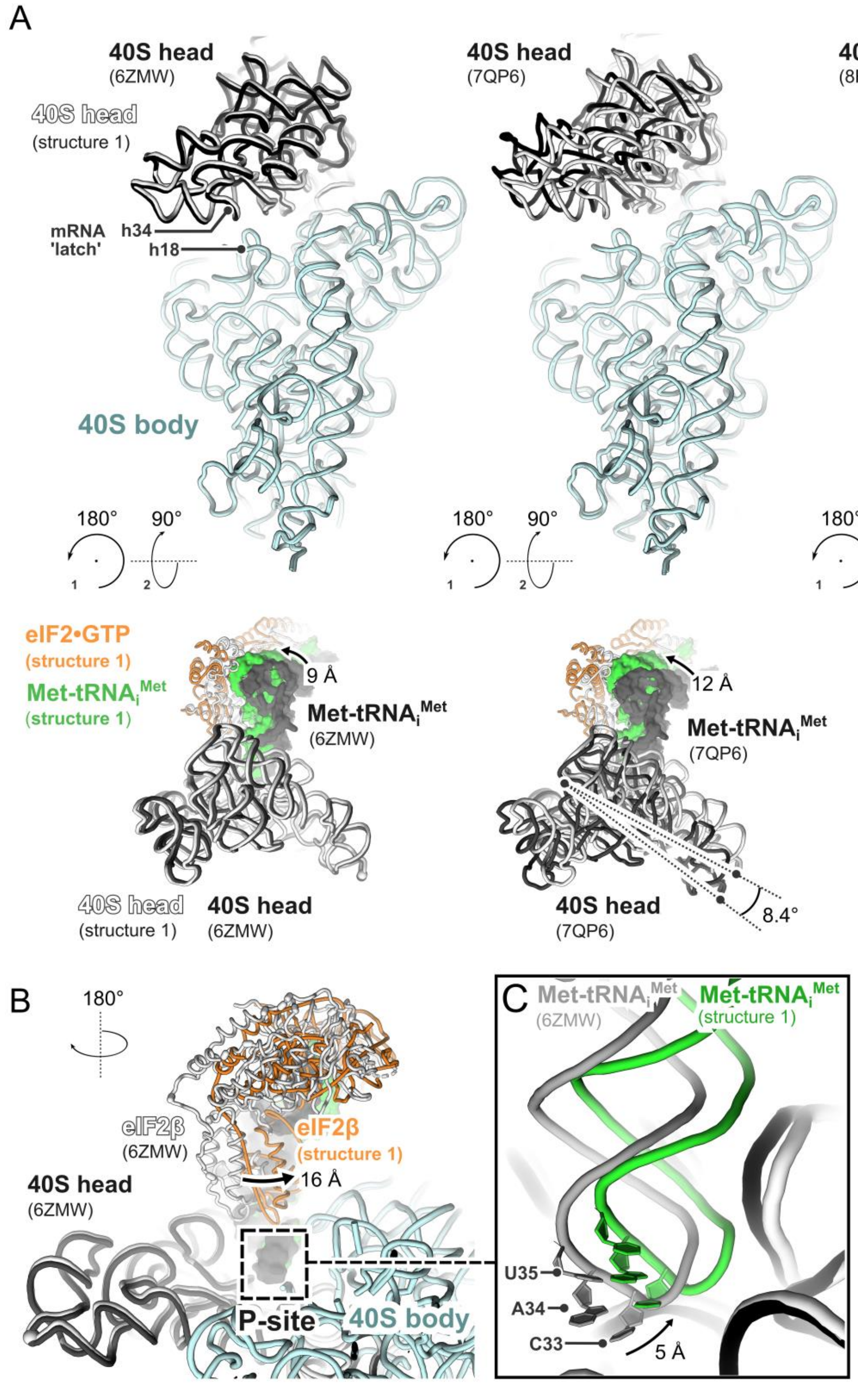

**Appendix Figure S6 – Comparison of 'open' 48S (structure 1) to other open/scanning human 48S structures**

**A,** Comparison of 'open' 48S complexes (this work, **structure 1**) with other mammalian open or scanning 48S structures, in two orthogonal views. In each panel, the position of the 18S rRNA comprising the 40S head (light grey), eIF2 (orange) and tRNA$_i$ (green) is shown relative to Data ref: PDB 8PJ1, 2024, Data ref: PDB 7QP6, 2022 and Data ref: PDB 6ZMW, 2020 (as indicated, dark grey) following structural superposition of the 40S body (light blue). In all cases, our eIF2 and initiator tRNA are tilted towards the E-site. Arrows indicate the relative displacement in the tRNA acceptor arm, measured at C64. Angles indicate the relative 40S head rotation, measured between G1680 in the hinge and G1285 at the tip of the 40S beak.

**B,** Rotated view, showing the relative position of the eIF2β subunit in the 'open' 48S **structure 1** (orange), compared to human scanning 48S Data ref: PDB 6ZMW, 2020.

**C**, Close-up view of the P-site, showing details of tRNA positioning compared to human scanning 48S Data ref: PDB 6ZMW, 2020. The anticodon loop of our $P_{OUT}$ initiator tRNA (green) is displaced by 5 Å towards the E-site (measured at C33).

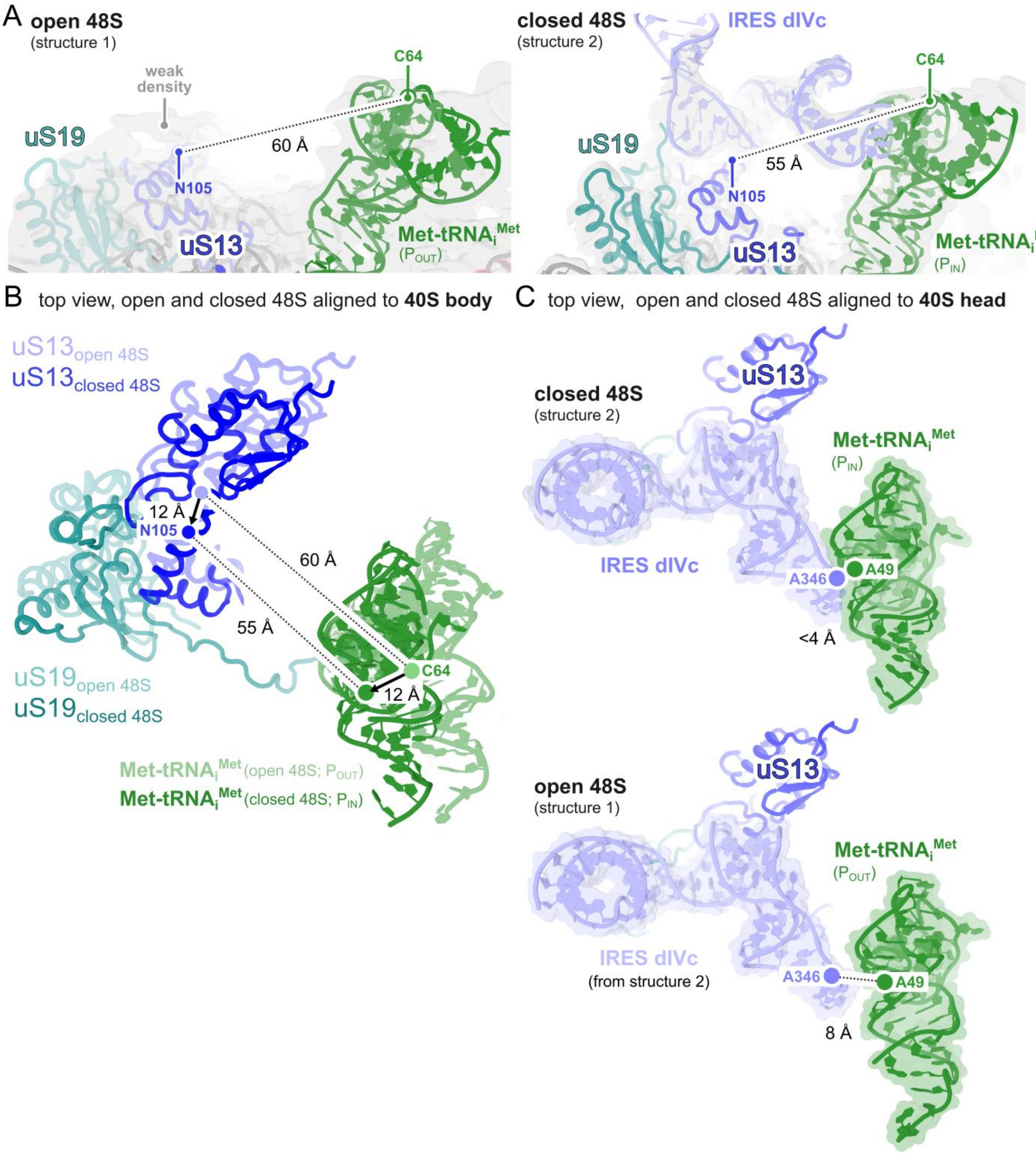

**Appendix Figure S7 – Analysis of geometry at the IRES binding site in 'open' and 'closed' 48S complexes**

**A,** Comparison of distances between uS13 residue N105 (dark blue) and tRNA$_i$ residue C64 (green) in 'open' / tRNA$_i$ P$_{OUT}$ (**structure 1**) and 'closed' / tRNA$_i$ P$_{IN}$ 48S (**structure 2**) complexes. Local-resolution weighted cryo-EM maps are shown (grey, contoured at 2.5 σ). Additional weak density adjacent to uS13 is indicated.

**B,** Superposition of **structures 1** and **2** (aligned to the 40S body) illustrate conformational changes at the IRES dIVc binding site associated with 40S head closure. Arrows indicate altered relative movements of uS13 (blue), uS19 (turquoise) and tRNAi (green). Measurements are between uS13 residue N105 and tRNA$_i$ residue C64, as above.

**C,** Superposition of **structures 1** and **2** (aligned to the 40S head) illustrate that the IRES dIVc GNRA-tRNA$_i$ interaction observed in 'closed' 48S complexes may not exist in 'open' 48S complexes, in which the tetraloop binding site is ~4 Å further away. Distances between IRES dIVc residue A346 (lilac) and tRNAi residue A49 (green) are indicated.

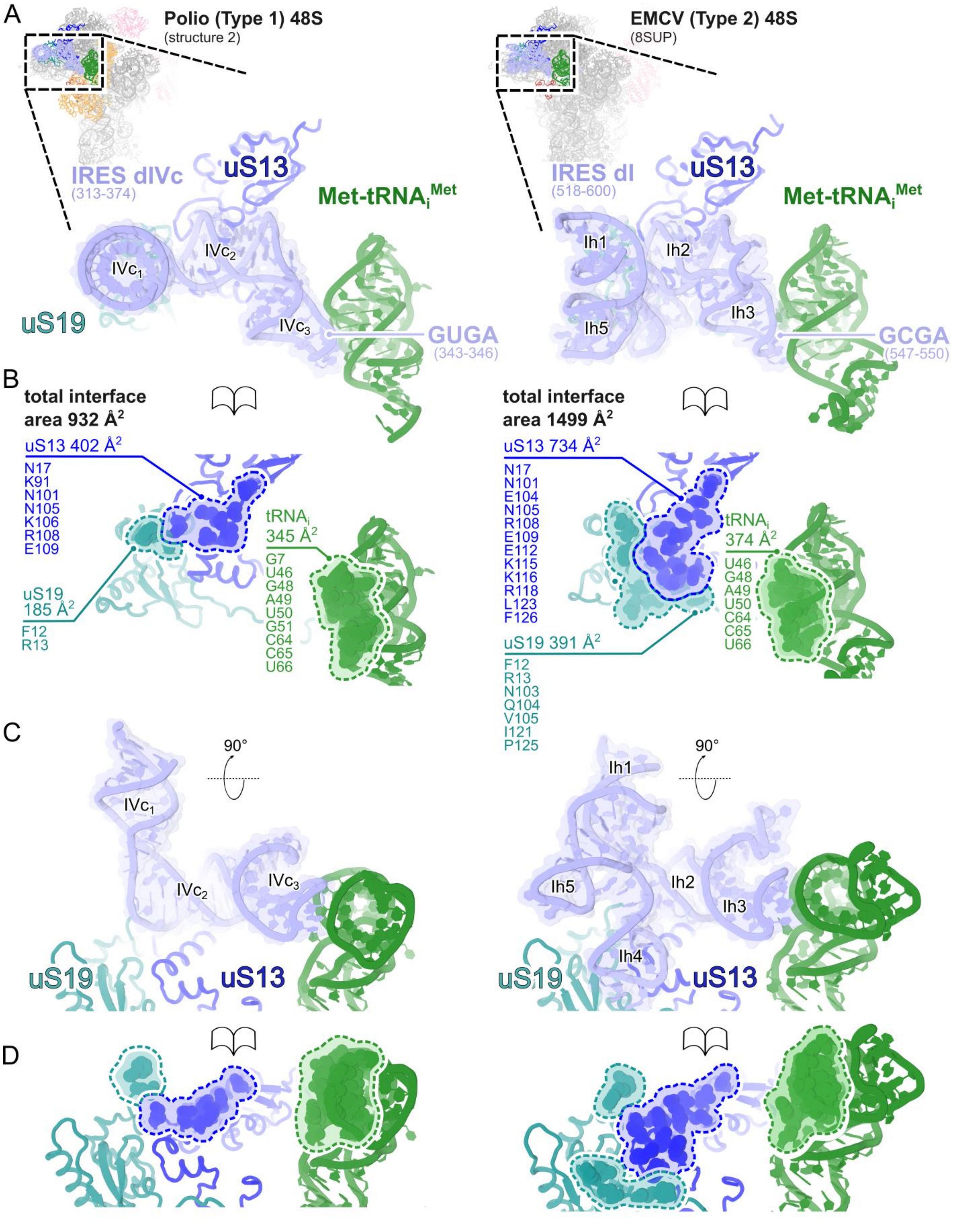

**Appendix Figure S8 – Comparison of 48S complex structures on Polio (Type 1) and EMCV (Type 2) IRESs**

**A,** IRES domains from Poliovirus (Type 1, **structure 2**) and EMCV (Type 2, Data ref: PDB 8SUP, 2026) bind to the 40S head at the same site, in a functionally equivalent manner during AUG recognition. Close-up top views show details of

IRES RNA (lilac) observed in ‘closed’ 48S complexes at $AUG_{586}$ (PV, *left*) and $AUG_{834}$ (EMCV, *right*). The GNRA-$tRNA_i$ interaction is conserved.

**B,** Peeled-apart view illustrating the IRES binding site comprising uS13 (blue), uS19 (turquoise) and $tRNA_i$ (green). Residues < 4 Å from the IRES RNA are listed and shown as spheres, and the calculated surface area (PDB-ePISA) for each interaction is reported.

**C-D,** As above (orthogonal view).

**Appendix Table S1 - Cryo-EM data collection and refinement statistics**

| | **Structure 1** 'open' 48S | **Structure 2** 'closed' 48S at $AUG_{586}$ | **Structure 3** 'closed' 48S at $AUG_{586}$ $dIVc_1$ position 1 | **Structure 4** 'closed' 48S at $AUG_{586}$ $dIVc_1$ position 2 |
|---|---|---|---|---|
| | PDB 32OB EMD-59031 | PDB 32OC EMD-59032 | PDB 32OE EMD-59033 | PDB 32OG EMD-59034 |
| **Data collection and processing** | | | | |
| Magnification | 105k | 105k | 105k | 105k |
| Voltage(kV) | 300 | 300 | 300 | 300 |
| Electron exposure (e-/Å$^2$) | 50 | 50 | 50 | 50 |
| Defocus range (μm) | -2.5 to -1 | -2.5 to -1 | -2.5 to -1 | -2.5 to -1 |
| Pixel Size (Å) | 0.831 | 0.831 | 0.831 | 0.831 |
| Symmetry imposed | C1 | C1 | C1 | C1 |
| Final particle images(no.) | 10,518 | 44,492 | 26,369 | 12,288 |
| Map resolution (Å) | 4.0 | 3.0 | 3.1 | 3.7 |
| FSC Threshold | 0.143 | 0.143 | 0.143 | 0.143 |
| Map resolution range (Å) | 3.1-8.1 | 2.6-12.7 | 2.7-12.1 | 3.0-15.7 |
| **Refinement** | | | | |
| Model Resolution (Å) | 4.0 | 3.0 | 3.1 | 3.7 |
| FSC threshold | 0.143 | 0.143 | 0.143 | 0.143 |
| Map sharpening B factor (Å$^2$) | -78.4 | -65.8 | -57.8 | -62.2 |
| Model composition | | | | |
| Non hydrogen atoms | 103,800 | 104,271 | 82,332 | 82,322 |
| Protein residues | 9022 | 9073 | 5690 | 5690 |
| Nucleotides | 1794 | 1797 | 1797 | 1797 |
| Ligands: $Mg^{2+}$, $Zn^{2+}$ | 4 | 3 | 3 | 3 |
| B factors (Å$^2$) (min/max/mean) | | | | |
| Protein | 22.0/618.1/217.8 | 8.8/538.8/221.0 | 9.6/405.6/102.42 | 3.4/379.9/95.0 |
| Nucleotide | 7.0/484.5/124.0 | 5.7/443.1/81.7 | 5.5/387.3/81.6 | 0.67/381.2/75.8 |
| Ligand | 56.3/333.9/182.9 | 22.9/151.6/70.0 | 33.1/195.3/88.0 | 26.2/189.4/81.9 |
| R.m.s deviations | | | | |
| Bond lengths(Å) | 0.004 | 0.005 | 0.003 | 0.003 |
| Bond angles (°) | 1.131 | 1.246 | 1.081 | 0.971 |
| Validation | | | | |
| Mol Probity score | 1.51 | 1.61 | 1.53 | 1.41 |
| Clashscore | 3.51 | 3.97 | 2.89 | 2.23 |
| Poor rotamers (%) | 0.66 | 1.47 | 1.54 | 1.20 |
| Ramachandran plot | | | | |
| Favoured (%) | 94.57 | 95.71 | 95.56 | 95.04 |
| Allowed (%) | 5.25 | 4.19 | 4.32 | 4.85 |
| Disallowed (%) | 0.20 | 0.10 | 0.12 | 0.11 |
